# Antibodies set boundaries limiting microbial metabolite penetration and the resultant mammalian host response

**DOI:** 10.1101/193581

**Authors:** Yasuhiro Uchimura, Tobias Fuhrer, Hai Li, Melissa A. Lawson, Michael Zimmermann, Mercedes Gomez de Agüero, Bahtiyar Yilmaz, Francesca Ronchi, Marcel Sorribas, Siegfried Hapfelmeier, Stephanie C. Ganal-Vonarburg, Kathy D. McCoy, Uwe Sauer, Andrew J. Macpherson

## Abstract

Although the mammalian microbiota is well-contained within the intestine and on other body surfaces, it profoundly shapes development and metabolism of almost every host organ, presumably through pervasive microbial metabolite penetration. The challenge is that most metabolites can be of both host and microbial origin. We developed a model to distinguish between microbial and host metabolites by stable isotope tracing using fully ^13^C-labelled live non-replicating *Escherichia coli*, differentiating ^12^C and ^13^C isotopes with high-resolution mass spectrometry. Hundreds of microbial compounds penetrated across 23 host tissues and fluids after intestinal exposure: subsequent ^12^C host metabolome signatures included lipidemia, reduced glycolysis and inflammation. Mucosal barrier maturation with transient microbial exposure increased early clearance of penetrant bacterial metabolites from the small intestine into the urine, independently of antibody induction. Induced antibodies curtailed microbial metabolite exposure at the intestinal surface, by accelerating intestinal bacterial transit into the colon where metabolite transport mechanisms are limiting.

## INTRODUCTION

The relationship between mammals and their resident microbiota shapes both the microbial communities and every host organ system from conception until death. The all-embracing scope of these effects is clear from comparing the differences between the same animal strain in the germ-free or colonised state (Smith et al., 2007). A few of these differences, such as natural killer T cell responses in the intestinal mucosa, depend on colonisation at critical developmental checkpoints, although most are rapidly reversible within weeks if a germ-free animal is colonised at any age, including intestinal secretory immunoglobulin (Ig)A induction (Gensollen et al., 2016). Given these ubiquitous interactions over all body systems, it is unsurprising that almost every disease model has been reported to be sensitive to microbial colonisation status (Blumberg and Powrie, 2012; Belkaid and Harrison, 2017; Sommer and Backhed, 2013).

Since the biomass of the microbiota is well separated from host tissues except during pathogenic infections, at first sight it is paradoxical that the presence of the microbiota shapes the cellular composition and function of non-intestinal mammalian organs so extensively, even when they are almost always sterile. The answer probably lies mainly in the penetration of microbial metabolites, which in some cases have been shown to exert powerful effects on host cell differentiation and metabolism. Microbial metabolite penetration of host tissues was first shown in principle by detecting aromatic compounds in colonised but not germ-free animals over 40 years ago (Cotran et al., 1960; Dayman and Jepson, 1969; Adamson et al., 1970; Scheline and Midtvedt, 1970; Peppercorn and Goldman, 1972). Typically, metabolites are inferred to originate directly or indirectly from microbial metabolism when they are present in host tissues only when animals are fully colonised or harbour specific compositions of microbial consortia (Wikoff et al., 2009; Williams et al., 2002; Nicholls et al., 2003; El Aidy et al., 2013). This approach is ambiguous for those compounds that can be synthesised either by host or microbial metabolism, because an increased metabolite level after microbial colonisation may be due to the propagation from microbes, or the host metabolic response to their presence.

The only alternative to robustly differentiate microbial or host origins of chemically identical metabolites are isotope tracing experiments. Given the complexity of metabolic conversions in the mammalian gut and host tissues, however, such stable isotope tracing has so far been used to follow gut microbial metabolism in an *in vitro* model where host effects were eliminated (de Graaf et al., 2010), or to show selective utilisation of carbon sources resistant to host digestion by *in vivo* incorporation into microbial RNA of a limited number of species within the intestinal consortia (Tannock et al., 2014). These labelled nutrient supplementation approaches are thus limited to spotlighting the fate of individual nutrients within the microbial communities themselves. Here we wanted to achieve a comprehensive view of the potential extent of metabolite penetration from the microbiota across a wide range of host tissues in a live animal, yet simultaneously to detect and distinguish the host metabolite response in each tissue as a result of gut colonisation.

Assessing the overall penetration of host tissues with microbial metabolites requires selective labelling of all microbial (but not host) molecules, yet avoiding exponential dilution of the label as bacteria use host or nutritional carbon sources to divide *in vivo*. For this purpose, we developed a strategy where germ-free mice are gavaged with a dose of either ^12^C-or fully ^13^C-labelled live *E.coli*. Since the employed HA107 strain cannot replicate in the germ-free intestine (Hapfelmeier et al., 2010), the ^13^C-label is not immediately and exponentially diluted *in vivo* through bacterial replication using host or food-derived ^12^C nutrients. High-resolution mass spectrometry that differentiates between ^12^C and ^13^C isotopes (Sevin et al., 2017; Fuhrer et al., 2017) was then used to reveal both the ^13^C host penetration by microbial metabolites and the ^12^C host metabolome response to the presence of the microbes. Although *E.coli* is a minor constituent of the adult human microbiota, it is a prominent member of the early life microbiota at a time when diversity is limited (Backhed et al., 2015), and thus models some aspects of early-life host-microbial molecular interactions from a low diversity microbiota. This parallel strategy enabled the detailed metabolic interactions in an unprecedented range of 23 different host tissues and fluids to be visualised at the level of individual molecules, concurrently showing microbiota molecular penetration and the resulting host molecular response.

There is a second problem in resolving the *in vivo* mechanisms that determine the molecular exchange between intestinal microbes and their mammalian host. The intestinal mucosa of germ-free animals, including its epithelial barrier, is extensively reprogrammed by exposure to intestinal microbes. The reprogramming encompasses epithelial cell differentiation, secretion of antibodies, antibacterial polypeptides, microvascular development and xenobiotic metabolic pathways (Hooper et al., 2001; Macpherson et al., 2000; Stappenbeck et al., 2002; Vaishnava et al., 2011; Gomez de Aguero et al., 2016). This means that microbes instruct the very mechanisms that build the dynamic interface separating the host, its intestinal microbes and their metabolites. Simply to challenge germ-free mice with ^13^C-labelled bacteria could overestimate molecular exchange by neglecting the immaturity of the barrier, yet to challenge a colonised animal would confound the distinction of microbial and host compounds based on the observed isotopes because of the much larger load of natural ^12^C metabolites from the resident microbiota. We have resolved this problem using a system that allows maturation of the intestinal barrier in germ-free animals through transient intestinal live microbial preconditioning without permanent colonisation (Hapfelmeier et al., 2010). This approach allowed us to compare microbial-host metabolite exchange directly in germ-free mice with or without prior microbial-driven mucosal adaptations. The combination of this new isotope tracing approach and the unique transient colonisation model has provided a unique insight into how microbial transit through the small intestine and propagation of microbial metabolites throughout the host are checked by secretory and metabolic adaptations of the intestinal mucosa.

## RESULTS

### The broad scope of metabolite penetration to systemic organs from intestinal microbes

Whilst a few microbial metabolites, such as short chain fatty acids or aryl hydrocarbon receptor ligands, are well-known to shape host metabolism and immune functions (Arpaia et al., 2013; Gomez de Aguero et al., 2016; Perry et al., 2016; Smith et al., 2013b), we do not know the full extent of microbial metabolite penetration, nor the entire scope of consequent host metabolic responses. To systematically assess the extent of host-microbial mutualism in metabolic terms, we set out to broadly identify the propagation of gut microbial metabolites to 16 mouse tissues and 7 fluids. For this purpose, we gavaged fully ^13^C-labelled or ^12^C live *E. coli* HA107, which cannot replicate *in vivo,* into C57BL/6 wild-type germ-free mice (Figure S1A). To achieve the required breath and depth of molecular discrimination, we employed high-resolution untargeted mass-spectrometry with high throughput methodology (Fuhrer et al., 2011; Sevin et al., 2017; Fuhrer et al., 2017). This could resolve over 1000 annotated compounds (M_r_ 50 – 1000 Da at 1mD resolution) and thus determine the mass shift showing the proportion of ^13^C or ^12^C within each molecule (Figures S1B and S1C). When the mice were challenged with bacteria whose molecules were verified as fully ^13^C-labelled after growth on a ^13^C-carbon source (Figures S1D and S1E) this showed bacterial metabolite penetration (Figures S1B and S1F). However, if the molecule concerned came from the ^12^C host response rather than the bacteria (whether or not the challenge bacteria were ^13^C-labelled) it would give a signal with the appropriate mass/charge ratio, but without the ^13^C mass shift (Figure S1C, illustrated for a hypothetical C_4_ compound).

As a starting point we investigated the expected propagation of microbial compounds from the intestine into the host. Two hours after gavage, a wide range of ^13^C-labelled bacterial metabolites had already reached almost all fluids and organs (Figures 1, 2A-C and S2A-S2D). However, the distribution of different compound classes varied. Bacterial amino acids, including essential amino acids that cannot be synthesised by host metabolism, penetrated very widely over all host tissues (Figures 1 and S2D). Many bacterial metabolites, including five and six-membered ring and heterocyclic compounds, were present in the serum and rapidly excreted in the urine (Figures 2B and S2C), but urinary lipid levels were generally low, consistent with known lipoprotein exclusion from glomerular filtration (Figures 2B and S2B). Bacterial fatty acids and glycerolipids were present mainly in the small intestinal tissue and mesenteric lymph nodes (Figure 2C) consistent with lymphatic uptake. Most compound classes apart from lipids were able to penetrate the peritoneal fluid of these naïve germ-free mice (compare analysis of different compound classes from Figure 2A shown in Figures S2A, S2C and S2D, with S2B).

**Figure 1.**
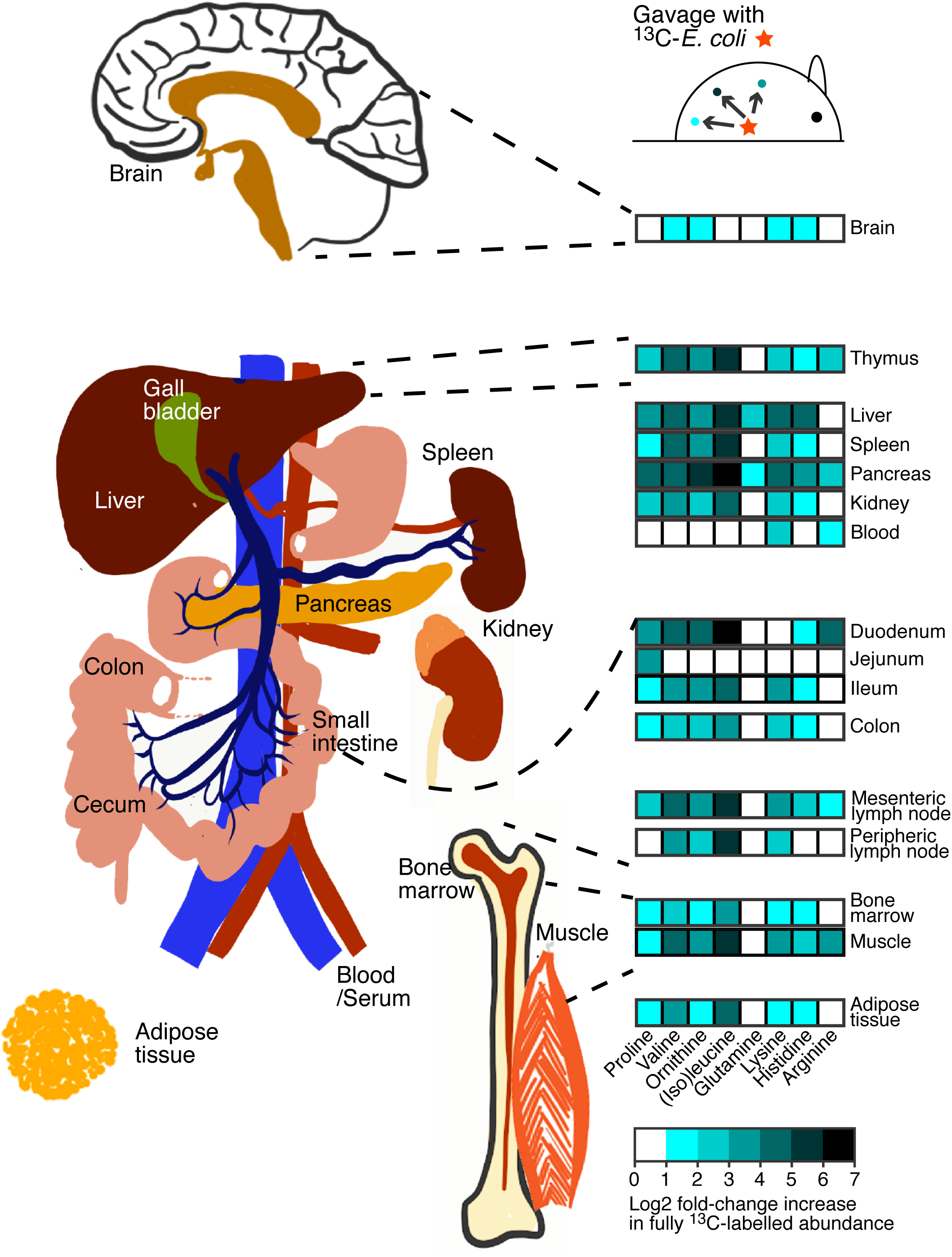
Depth of penetration of ^13^C-labelled bacterial amino acids from the intestine to systemic sites of the body. Germ-free C57BL/6 mice were gavaged with ^13^C- or ^12^C-labelled HA107 or PBS as control (n = 3 - 4 mice per group [Figure S1A]). Differential ions detected in different host tissues after 2 hours annotated as fully^13^C-labelled amino acids are indicated where the identical mass did not exhibit any significant change in the ^12^C to PBS comparison ([Figure S1B and S1C] positive Log -fold change cutoff of 1 and p-value < 0.05). Results are representative of three experiments.

**Figure 2.**
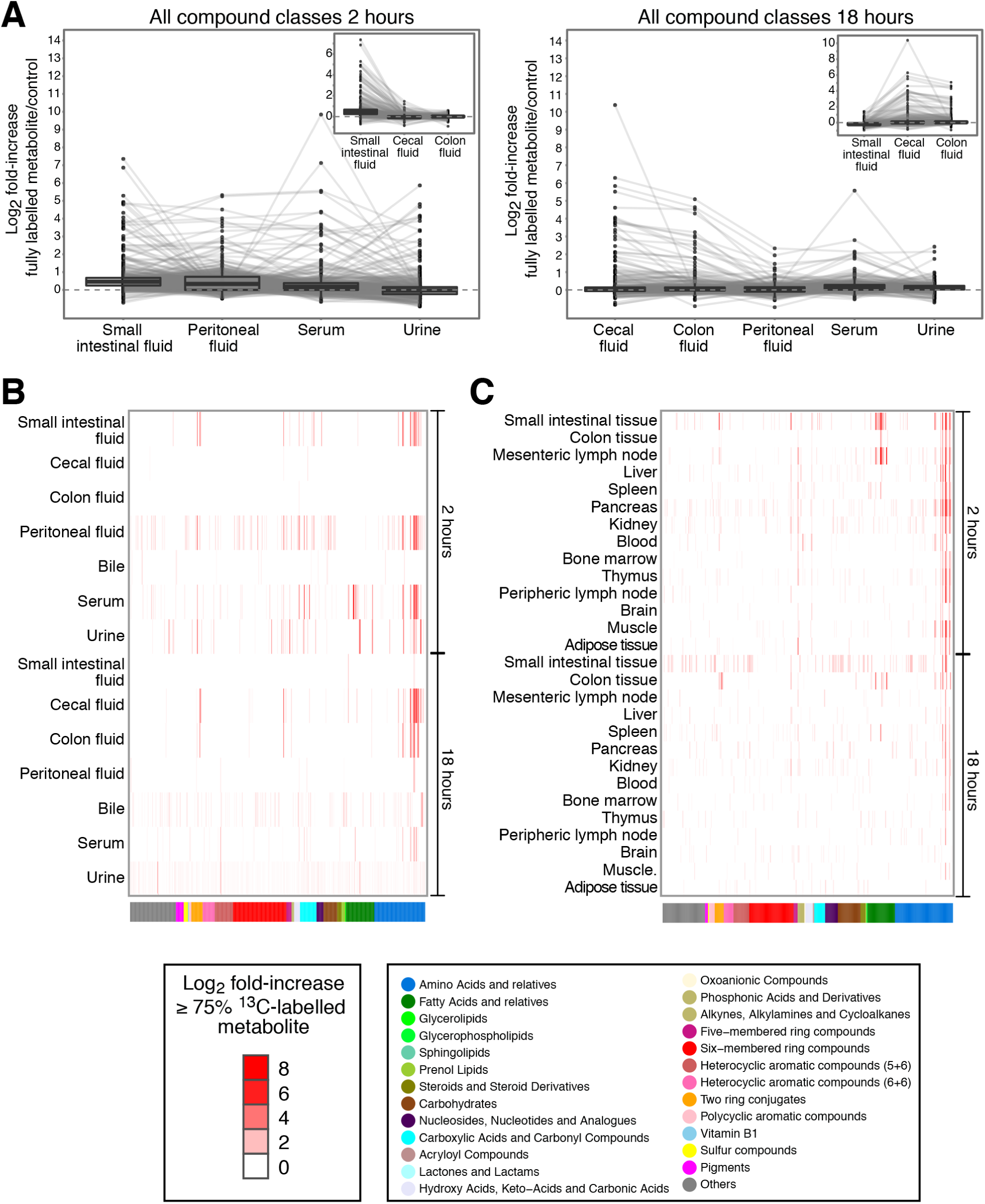
Penetration of ^13^C-labelled bacterial metabolites from intestine to host tissues and fluids. Germ-free C57BL/6 mice were gavaged with ^13^C-labelled HA107 or PBS as control (n = 3 - 4 mice per group). Fluid samples were collected 2 or 18 hours later and analyzed with Q-TOF mass spectrometry. All ions which could be annotated as metabolite isotopes with ≥75% of the carbon backbone being ^13^C labelled were plotted. **(A)** Left panel. Samples from small intestinal fluids, peritoneal fluid, serum and urine at 2 hours are shown, with lines connecting identical metabolites. The inset shows intestinal samples from small intestinal fluids, cecal fluids and colon fluids also at 2 hours. Right panel shows 18 hour cecal, colon, peritoneal, serum and urine fluid samples with (inset) small intestinal, cecum and colon fluids. **(B)** Heat map showing cluster analysis of all ^13^C-labelled annotated metabolites (p < 0.05, ≥ 3 mice/group) in different host fluids at 2 hours and 18 hours after gavage. Metabolite classes are shown in the lower colour bar and legend. Results are representative of three experiments**. (C)** As for B, except ^13^C-labelled annotated metabolites in host tissues are shown.

By 18 hours, the front of detected ^13^C-labelled metabolites had moved along the intestine to reach the cecum and the colon (compare Figure 2A insets and S2A-S2H insets at 2 and 18 hours) consistent with the known murine small and large intestinal transit times (Padmanabhan et al., 2013). Although bacterial metabolites had been cleared from the small intestinal fluid and peritoneum by 18 hours, they were still present in small intestinal tissues and being excreted at low levels in the urine and the bile (Figures 2B, 2C and S2E-S2H).

Since the HA107 system depends on cell wall auxotrophy to abrogate *in vivo* replication, we asked whether increased bacterial fragility of the HA107 system generates increased systemic exposure of bacterial metabolites. To address this, we compared penetration of compounds from HA107 or its non-auxotrophic parent strain. In this case, bacterial compounds were labelled by growth in ^14^C glucose and bulk transfer of radioactivity was estimated. The use of ^14^C metabolic labelling shows only the bacterial origin of the radiolabel but is uninformative about the original molecular identity of the bacterial compound. However, for this experiment it had the advantage that we could sensitively measure overall ^14^C transfer, whereas bacterial replication *in vivo* would dilute the proportion of ^13^C-label leading to a loss of sensitivity measured on a per molecule basis, and potential underestimation of penetration. We found overlapping ranges of radiolabel recovery in serum, urine and host tissues and no significant differences at most timepoints, regardless of whether or not the test bacteria were engineered with cell wall auxotrophy (Figures 3A, 3B and S3A-C). Hence there is no evidence that potential fragility of the employed HA107 strain would liberate significantly more metabolites than wild-type *E.coli*.

**Figure 3.**
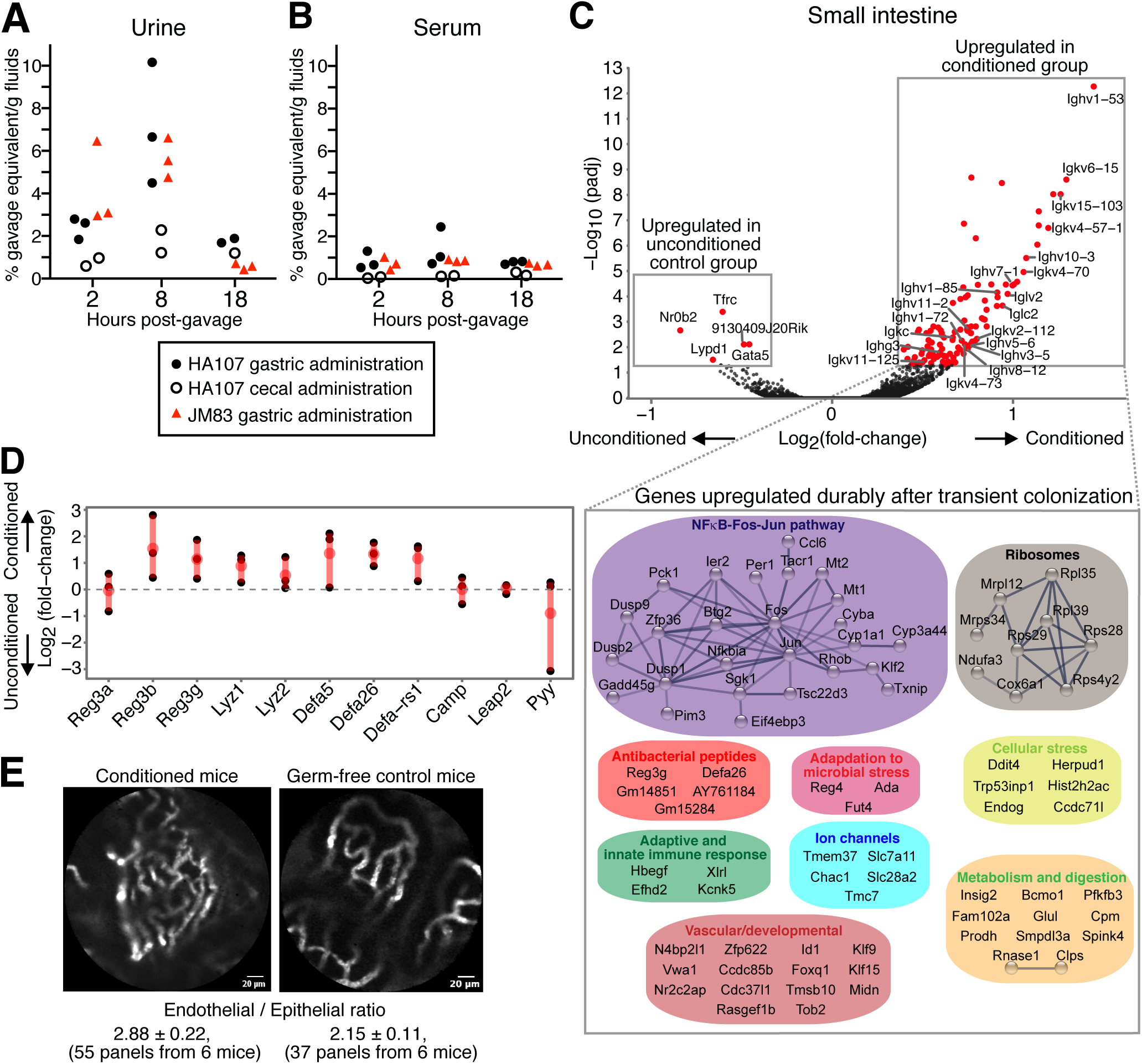
Kinetics of metabolite penetration and durable intestinal conditioning responses to E.coli HA107. **(A-B)** ^14^C-labelled E. coli HA107 (after growth on ^14^C-glucose; closed circles) was administered directly into the stomach or mixed directly with the cecal contents (open circles) of germ-free C57BL/6 mice. For comparison, ^14^C-labelled *E. coli* JM83 (the replication competent parent strain of HA107; red triangles) was administered into the stomach of a separate group of mice. ^14^C-radioactivity in the urine **(A)** or serum **(B)** was monitored at 2, 8, and 18 hours later. Results are expressed as a percentage of the gavage dose/g fluid after subtraction of background. **(C)** RNAseq analysis of small intestinal tissue from C57BL/6 germ-free mice that had been primed with HA107 transitory colonisation two weeks previously, compared with small intestinal gene expression from control germ-free mice. Upper panel: volcano plot shows significantly altered gene expression (padj < 0.05, n≥3) in HA107-preconditioned germ-free mice or in sham-treated germ-free mice, shown on the right and left of the x-axis respectively. Immunoglobulin genes are annotated. The expanded box shows a STRING-DB representation of significantly upregulated non-immunoglobulin genes in preconditioned mice. **(D)** Expression of transcripts coding microbiocidal peptides [inline]. **(E)** Live intestinal microvascular endomicroscopy imaging. 70 kDa FITC-dextran was injected intravenously into HA107-preconditioned C57BL/6 germ-free and control germ-free mice. The FITC-dextran fluorescence was visulized in the small intestinal villus microvasculature with live Cellvizio endomicroscopy. The [inline] of the endothelial/epithelial ratio of each group (n=6) is indicated below the respective panels (t-test p < 0.02).

We concluded that there is extensive penetration of an extremely broad range of bacterial-derived molecules into host tissues from the germ-free intestine, even though live bacteria penetrate only as far as the local (mesenteric) lymph nodes (Hapfelmeier et al., 2010). These results did not distinguish the relative penetration through the small or large intestinal surfaces. Furthermore, the germ-free intestine is not adapted to the presence of intestinal microbes, for example through the induction of secretory IgA or IgM, antibacterial peptides and a consolidated barrier function (Vaishnava et al., 2011; Salzman et al., 2010; Petersson et al., 2011; Hapfelmeier et al., 2010). We therefore needed to address how these different aspects of prior mucosal adaptation could shape microbial-host molecular exchange at both small and large intestinal surfaces.

### Preferential penetration of molecules from intestinal microbes through the small intestine

We also asked through which gut compartment the microbial metabolites primarily penetrate? The small intestine has minimal mucus coverage and is highly adapted for molecular uptake, whereas the large intestine has a thick layer of protective mucus overlying the epithelium (Johansson et al., 2008). We therefore expected that the most susceptible surface to microbial metabolite exchange would be the small intestinal mucosa. This was confirmed experimentally by comparing uptake of ^14^C HA107 after delivery into the stomach or after direct *in vivo* mixing with the fluid caecal contents in a separate group of germ-free mice. Labelling with ^14^C was also used in this experiment to detect host exposure to microbial metabolites with maximum sensitivity, even if they underwent secondary metabolism in the host. In contrast to wide dissemination of radiolabel from the upper intestine, hardly any radiolabel was transferred when the bacteria were directly mixed with the cecal contents (Figures 3A, 3B and S3A-C). The highest recovery of radiolabel that had originated in the bacteria was in the urine (Figure 3A). This verified that the small intestine was the main absorptive surface for extensive microbial molecular exchange with host organ systems and verified results with the specific molecular ^13^C system that the urine is a highly effective immediate elimination route.

The lower small intestine is normally colonised with intestinal microbes, so microbial molecular exposure has potentially important biomedical consequences. Mice are naturally coprophagic, but even adult humans or infants with reduced intestinal motility can have high densities of microbes in the proximal small intestine (Donowitz et al., 2016; Jones et al., 2011). In the developing world, small intestinal microbes cause childhood environmental enteropathy – a very serious health problem characterised by intestinal inflammation, malnutrition, growth stunting and poor immune responses, affecting approximately 25% of children aged under 5 years (Korpe and Petri, 2012).

### Maturation of the intestinal mucosal barrier following transient live microbial exposure is durable and involves immune and non-immune transcriptional reprogramming

Environmental enteropathy that manifests as malnutrition refractory to caloric supplementation is associated with compensatory secretory IgA responses, directed against microbes responsible for the loss of intestinal function (Beatty et al., 1983; Kau et al., 2015). Furthermore, secretory IgA is known to be dominantly induced by small intestinal microbes (Bunker et al., 2015). We therefore wished to address whether antibodies or other barrier functions that are induced as a result of exposure to intestinal microbes constrain microbial metabolite penetration and how this impacts the body metabolome of the host response.

To assess the impact of mucosal protective factors against bacterial metabolite exchange in the intestine and its subsequent host effects, we needed a system that experimentally uncoupled maturation of the mucosa and its barrier functions from permanent colonisation. The HA107 system was potentially well suited for this. Even though the mice return to germ-free status approximately 48 hours after exposure, a persistent and specific secretory IgA response directed against the auxotrophic strain is induced (Hapfelmeier et al., 2010). However, determining the effects of the barrier on molecular penetration required that other non-immune, as well as immune barrier functions would also be durably induced by preconditioning with live transitory colonisation. This would then provide the tools to experimentally address how bacterial metabolite exposure is limited by rechallenging the germ-free mice with or without preconditioning (Figure S1Aii).

To examine the range of mucosal adaptations, we first conditioned germ-free mice through live reversible *E. coli* HA107 colonisation and carried out transcriptomic analysis of the intestinal mucosa 2 weeks later, once mice had returned to germ-free status. This showed an extensive range of very durable adaptations (Figure 3C and expanded box). In line with our earlier flow cytometry analysis and immunohistochemistry results that demonstrated a persistent IgA response (Hapfelmeier et al., 2010), we found that a series of transcripts representing Ig loci were significantly upregulated 2 weeks after experimental challenge with HA107, even though the mice had been verified germ-free for 12 days after live bacterial exposure. In addition, there was upregulation of genes with known functions in xenobiotic and metal metabolism, mitochondrial function, ion channels, angiogenesis (Figure 3C expanded box) and epithelial microbiocidal peptide expression (Figure 3D). In a complementary approach, *in vivo* endomicroscopy was used to compare the endothelial to epithelial ratios in the small intestine of treated and sham control animals (Figure 3E). This demonstrated that earlier observations of an increased microvascular density induced by permanent colonisation with Bacteroides thetaiotaomicron (Stappenbeck et al., 2002) were also shown to be durable after transitory colonisation 14 days earlier.

We concluded that even transitory mucosal conditioning by intestinal bacteria results in extensive re-programming of epithelial and stromal elements of the intestinal mucosa of adult animals, in addition to the known immune adapative response: in both cases this is durable after the mice have returned to germ-free status. This showed that we could evaluate the barriers to microbial molecular penetration through the intestine in an adult germ-free animal that had been preconditioned by HA107 exposure 14 days earlier, to experimentally mature its intestinal barrier. Since both conditioned and control-treated mice were germ-free at the time of challenge, we avoided competition effects from a resident intestinal microbiota and confounding contamination with ^12^C metabolites from resident microbes (schema in Figure S1Aii).

### Immunoglobulin induction accelerates small intestinal transit of bacteria and their metabolites

To determine the effect of mucosal adaptation after microbial exposure on the depth of metabolite penetration, and to discriminate the effects of antibody induction, we preconditioned germ-free C57BL/6 wild-type and J _H_^-/-^ mice. J _H_^-^deficient mice lack all antibody isotypes because of deletion of the J-segments of the immunoglobulin heavy chain locus (Chen et al., 1993). Administration of transient doses of intestinal HA107 results in a dominant secretory IgA response and minimal serum response in the intestinal mucosa in C57BL/6 wild-type mice (Hapfelmeier et al., 2010; Gomez de Aguero et al., 2016). This comparison between wild-type C57BL/6 animals and J_H_-deficient mice was used in order to avoid the confounding effects of compensatory intestinal IgM and overinduction of systemic IgG in animals selectively deficient for the IgA isotype (Harriman et al., 1999).

Groups of germ-free wild-type C57BL/6 and J _H_^-/-^ mice (≥18 per group) were conditioned with HA107: separate groups of each strain (≥18 per group) were treated in parallel with sterile PBS as controls. Preliminary experiments established that ^14^C HA107-treated mice had no residual detectable radiolabel in host fluids or tissues 14 days after treatment. Two weeks after the conditioned HA107-treated mice had returned to germ-free status, they were divided into 3 groups of ≥6 animals of each strain to be challenged with a single dose of ^13^C- or ^12^C-HA107, or sterile PBS, administered directly into the stomach (Figure S1Aii). To systematically investigate how antibody deficiency modulates the effect of preconditioning, we compared the relative difference of positive fold-changes due to propagation of ^13^C-labelled compounds between pre- and unconditioned germ-free mice from the C57BL/6 and J _H_^-/-^ strains (Figures S1B, S1C, 4A and 4B).

Results from the two different host strains (that were processed throughout at all timepoints in parallel) are colour-coded and shown alongside each other. Two hours after bacterial challenge, the conditioned C57BL/6 mice showed increased bacterial metabolites in the cecal fluid and colonic fluid compared with the antibody-deficient J _H_^-/-^ strain (p < 0.001 for all measured compounds and for different compound classes considered individually in colonic fluid: Figures 4A and 4B). In contrast, bacterial compounds were enriched in the small intestinal fluid of unconditioned mice compared to conditioned mice (Figure 4A). Since these effects were generalised across compound classes, we hypothesised that overall elevated levels of bacterial metabolites in the colonic fluid of antibody-expressing mice was due to induced secretory Ig accelerating the transit of intact intestinal bacteria, either through limiting bacterial motility and adherence, or exclusion effects resulting from increased size of the antibody-coated microbes.

**Figure 4.**
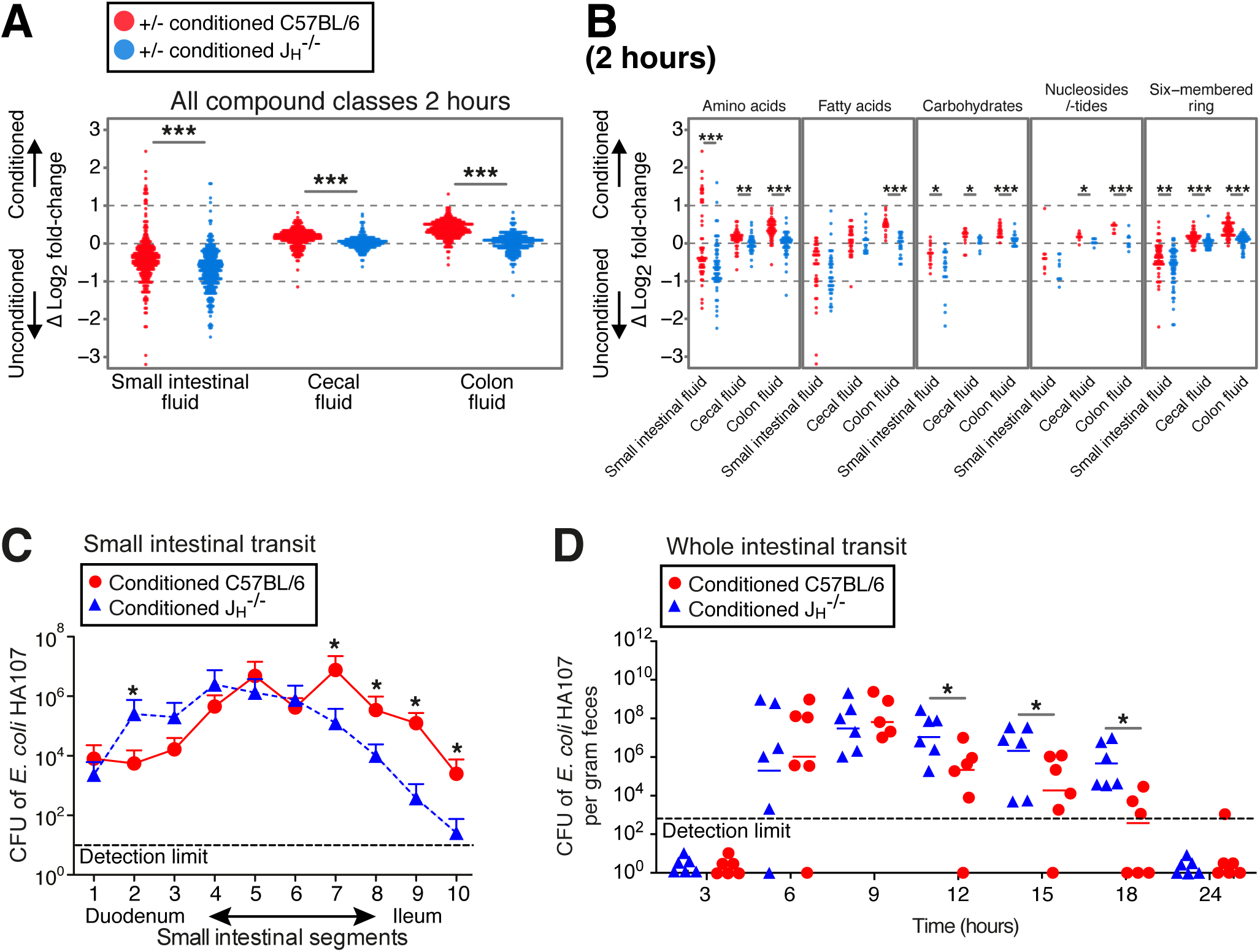
Transient intestinal microbial conditioning enhances clearance of bacteria and their metabolites from the intestine of C57BL/6 mice compared with antibody deficient J _H_^-/-^ mice. Germ-free C57BL/6 mice (red symbols) or antibody-deficient J _H_^-/-^ mice (blue) were preconditioned with HA107 or control-treated. Twelve days after all groups had returned to germ-free status they were gavaged with ^13^C-labelled HA107 or PBS as control (n = 3 to 4 per group) and Log_2_ fold-changes were calculated [Figure S1Aii]. (A) Samples from small intestinal fluid, cecal fluid and colon fluid were collected at 2 hours after HA107 oral gavage and analyzed with Q-TOF mass spectrometry. The difference between the Log_2_-fold values of preconditioned and unconditioned mice is shown for ions annotated as metabolite isotopes with ≥75% of the carbon backbone being ^13^C labelled. Fold-changes based on low abundant ions (counts < 200) in either ^13^C or PBS control samples were excluded. Mann-Whitney U test *** = p < 0.001. **(B)** As for panel A, but the plot shows individual compound classes. **(C)** Small intestinal transit of replication-deficient bacteria in wild-type mice and antibody-deficient mice. HA107 preconditioned C57BL/6 mice (red symbols) were compared with HA107 preconditioned antibody-deficient J _H_^-/-^ mice (blue) 2 hours after challenging with 10^7^ CFU HA107 delivered into the stomach. The whole small intestine was divided into 10 sections and the bacteria in contents were plated with auxtrophic supplements. Colony-forming units (CFU) of each section are shown [Inline]. (D) Whole intestinal transit of replication-deficient bacteria in wild-type mice and antibody-deficient mice. HA107 preconditioned C57BL/6 mice (red symbols) or antibody-deficient J_H_^-/-^ mice (blue) were gavaged with 10^8^ CFU HA107. Feces were collected after the indicated time after HA107 gavage and plated. The colony-forming units (CFU) are shown (* = p < 0.05).

Prior to directly testing the hypothesis that induced immunoglobulins accelerate the small intestinal transit of intestinal bacteria, we verified in our system that maximum levels of secretory IgA directed against intestinal bacteria were expressed in the small intestine (Bunker et al., 2015), even in mice that have returned to germ-free status (Figure S4A), and that these antibodies coat the HA107 challenge organisms (Figures S4B-S4D), confirming earlier results with bacterial flow cytometry (Hapfelmeier et al., 2010). Direct in vivo demonstration of accelerated intestinal bacterial transit was obtained by faster clearance of a test dose of HA107 from preconditioned germ-free C57BL/6 mice compared with preconditioned germ-free J _H_^-/-^ controls in two ways: first, by examining the trajectory of transit in different segments of the small intestine 2 hours after administration of the test dose into the stomach (Figure 4C) and secondly by following the kinetics of bacterial shedding in the feces after transit the through the entire intestine of the intact animal in vivo (Figure 4D). In both experiments, small and total intestinal transit was accelerated in the conditioned C57BL/6 wild-type, but not the conditioned J _H_^-/-^ strain. By 18 hours, most of the load of challenge bacteria had cleared from the colon of the C57BL/6 wild-type mice. Similar results were obtained comparing pretreated and non-treated control C57BL/6 mice (not shown). From this comparison between conditioned mouse strains that can, or cannot induce an antibody response, we concluded that the observed accelerated transit effect was due to induced antibodies independently of other non-immune mucosal conditioning effects. Therefore mucosal antibody induction and secretion limits the small intestinal dwell-time of intestinal bacteria and hence the exposure of the host to their small intestinal metabolites.

### Non-immune mechanisms of prior intestinal conditioning with live microbes determine the bodily distribution and excretion of microbial metabolites

Having shown that induced intestinal immunoglobulins accelerate intestinal bacterial clearance from the small intestine, we next wished to investigate how conditioning the intestinal mucosa would affect the distribution of microbial metabolites in different compartments of the body and excretion in the urine.

So far, we had found that without conditioning ^13^C bacterial metabolites appear in the peritoneal fluid of C57BL/6 mice early after the bacterial challenge (Figures 2A, 2B, S2A-S2D and 5A). This effect was also seen in unconditioned antibody-deficient mice (Figure 5A) and applied to nearly all compound classes (Figure S5A). However, accumulation in the peritoneum was abolished by preconditioning and instead there was preferential elimination in the urine: this effect was antibody-independent as it was seen in both wild-type and antibody-deficient animals (Figure 5A). At later times there was persistent uptake of bacterial molecules from the intestine into the serum of conditioned antibody-deficient mice (Figure 5B) reflecting the increased dwell time of bacteria in the small intestine (Figures 4C and 4D). Since metabolites are very effectively filtered into and concentrated within the urine, and the lower serum concentrations remain constant until at least 18 hours (compare Figures 3A and 3B), we concluded that the general preconditioning response improves early clearance of metabolites into the urine. Given that a non-immune effect of bacterial preconditioning is to increase the intestinal microvascular density (Figure 3E), the conditioned mucosa likely favours metabolite uptake through the intestinal microvasculature and rapid clearance into the urine at early times as an antibody independent effect (Figure 5A). Increased cardiac output and peripheral perfusion have also been observed by comparing germ-free and colonised mice, so it is likely that preconditioning also has hemodynamic effects that increase renal filtration and tubular excretion (Smith et al., 2007).

**Figure 5.**
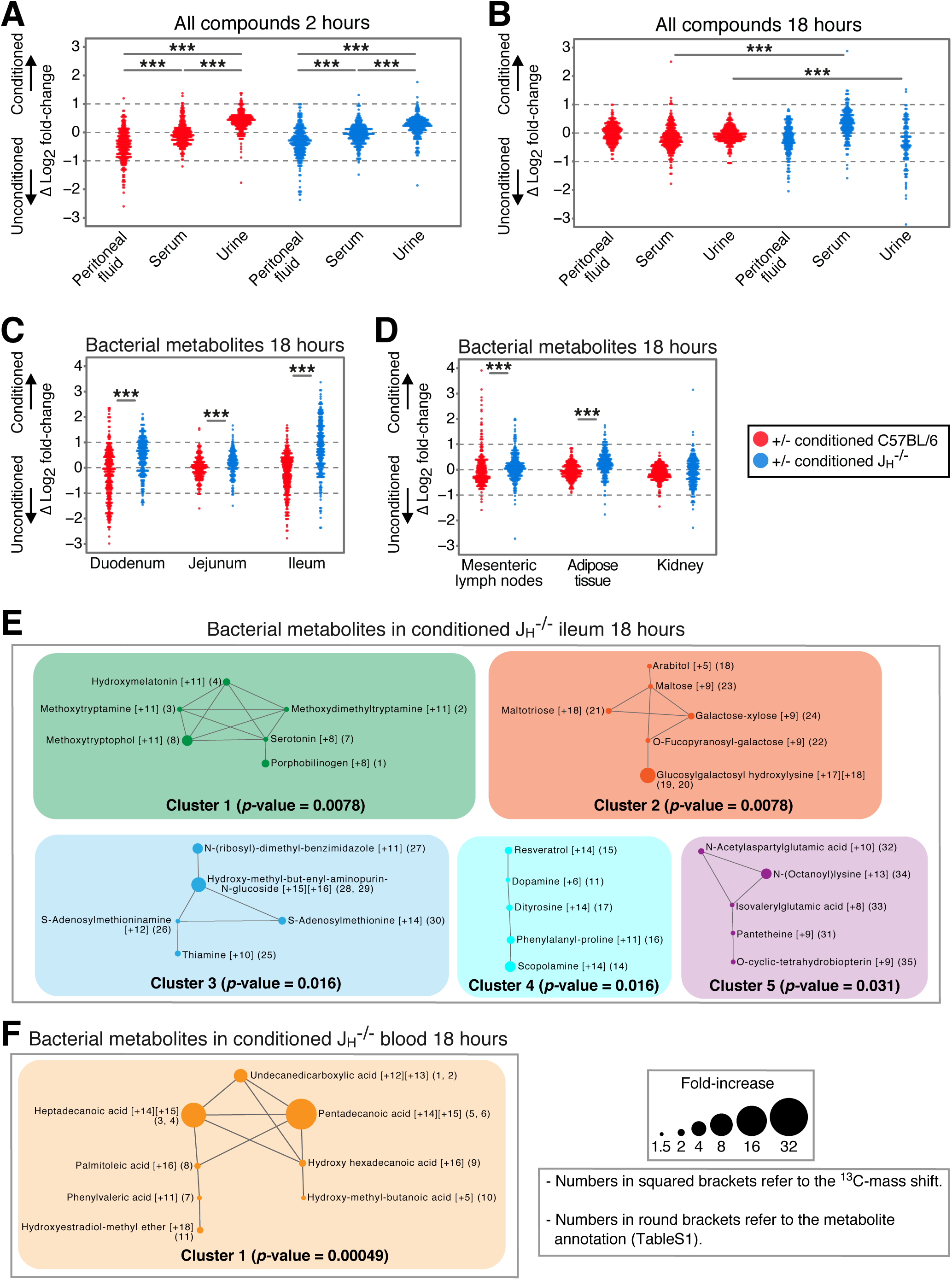
Transient intestinal microbial conditioning enhances antibody-independent early clearance of bacterial metabolites into the urine. Germ-free C57BL/6 mice (red symbols) or antibody-deficient J_H_^-/-^ mice (blue) were preconditioned with HA107 or control-treated, returning to germ-free status twelve days prior to re-challenge with a gastric dose of fully ^13^C-labelled or ^12^C-HA107 as in the legend to Figure 4. **(A-B)** Log_2_ fold differences between HA107-preconditioned and unconditioned germ-free mice for individual annotated ions representing metabolites containing ≥75% ^13^C are shown for peritoneal fluid, serum or urine 2 hours (A) or 18 hours (B) after the challenge dose. **(C-D)** As for panel B but showing persistence of bacterial metabolites in intestinal (C) or systemic (D) tissues of antibody deficient mice at 18 hours. **(E)** Over 1.5-fold increased bacterial metabolites (≥75% ^13^C) in ileal tissues at 18 hours of conditioned antibody-deficient mice were applied to Metamapp to calculate pairwise Tanimoto chemical similarity indices. Cytoscape software was used for network visualization. Numbers in squared brackets show the mass shift of each metabolite and numbers in round brackets correspond to listing for each metabolite in Table S1 also showing chemical annotations. Metabolites with Tanimoto chemical similarity ≥ 0.7 are interconnected. The size of nodes correlates with fold-increase. The paired Mann-Whitney U test was applied to calculate the p-value of each network. **(F)** As for panel E, except analysis of blood metabolite networks. Results are representative of three experiments.

Therefore, the non-immune effects of conditioning the intestinal mucosa are to increase the early clearance of penetrant bacterial metabolites from the small intestine into the urine and prevent accumulation in the peritoneal fluid. Antibody induction supplements this clearance effect by curtailing small intestinal mucosal exposure through accelerating intestinal bacterial transit out of the small intestine into the colon, where the permeability and metabolite transport mechanisms are limiting.

From the results showing increased dwell time of bacteria in the small intestine of antibody-deficient mice, we expected to detect increased bacterial metabolites in intestinal tissues. This was confirmed in the small intestine, mesenteric lymph nodes and some systemic tissues (Figures 5C and 5D). Specifically, we found that these bacterial metabolites in the ileum of conditioned antibody-deficient mice included potentially inflammatory biogenic amine indole derivates such as methoxytryptamine, methoxytryptophol, methoxydimethyltryptamine and serotonin (Figure 5E). In some cases, there was evidence of ^12^C incorporation into the dominantly ^13^C-labelled metabolites (Table S1 specifies molecular mass-shift results from Figure 5E), indicating formation after additional host or bacterial metabolism *in vivo*. There was also evidence of bacterial saccharides such as arabitol and galactose-xylose normally associated with bacterial but not host metabolism (Figure 5E). The serum of conditioned antibody-deficient mice contained increased bacterial non-esterified fatty acids, most of which were fully ^13^C-labelled (Figure 5F and Table S1). Bacterial-host metabolite crosstalk therefore includes direct bacterial lipid exposure with the potential to activate the host innate immune system and contribute to the systemic lipid load. To show increased exposure to lipids as a function of colonisation even in permanently colonised animals, we compared the cecal fluids of germ-free adult C57BL/6 mice and the same strain permanently colonised with a standardised stable microbiota of 12 defined organisms (Brugiroux et al., 2016), both fed with the identical sterilised chow diet (Figures S5B and S5C). It is therefore likely that the bacterial lipid load makes a significant contribution to host lipids, particularly in the context of defective intestinal barrier function.

### Persisting exposure to bacterial metabolites and increased host metabolomics response in antibody-deficient mice

Given the pervasive penetration of bacterial-derived metabolites in many different host tissues, including inflammatory triggers such as biogenic amines and lipids, especially when the immune barrier is compromised through defective antibody production (Figure 5B and 5C), a metabolic host response was expected. We questioned whether this host response would be mainly restricted to the intestine, or seen across all host tissues. To address this we exploited the discrimination power of the stable isotope tracing system and interrogated the molecular signatures by analysing the changes in ^12^C metabolite levels across different host tissues that were independent of whether a ^13^C or ^12^C bacterial challenge dose had been given (Figure S1C).

Tissue ^12^C-metabolite measurements 18 hours after a transient dose of either ^13^C-labelled or ^12^C HA107 showed both increases and loss of different specific metabolites in intestinal tissues (Figure 6A) and in systemic tissues (Figure 6B) of conditioned antibody-deficient compared with wild-type mice. The specific classes of host compounds that are altered at early or late times after transitory bacterial exposure are shown in Figure 6C. Since many of these compounds are currently not annotated in metabolic pathways, we used chemical similarity clustering as a surrogate for metabolic pathway analysis. For example, in the antibody-deficient ileal tissues, Tanimoto chemical similarity network analysis of > 2-fold changes showed increased host lysophospholipids, phosphatidic acids and release of non-esterified fatty acids and the phospholipid head-groups such as arachidonic acid (Figure 7A). This response was consistent with the 2.7- fold increased transcription of ileal phospholipase A2 (Leslie, 2015) that we observed after bacterial challenge in the intestine in the absence of induced secretory antibodies. An inflammatory response was also manifest by generation of downstream arachidonic acid-derived host production of prostaglandins and leukotrienes including leukotriene E3 and prostaglandins A4, E2, F1a and F2a (Figures S6A and 7A).

**Figure 6.**
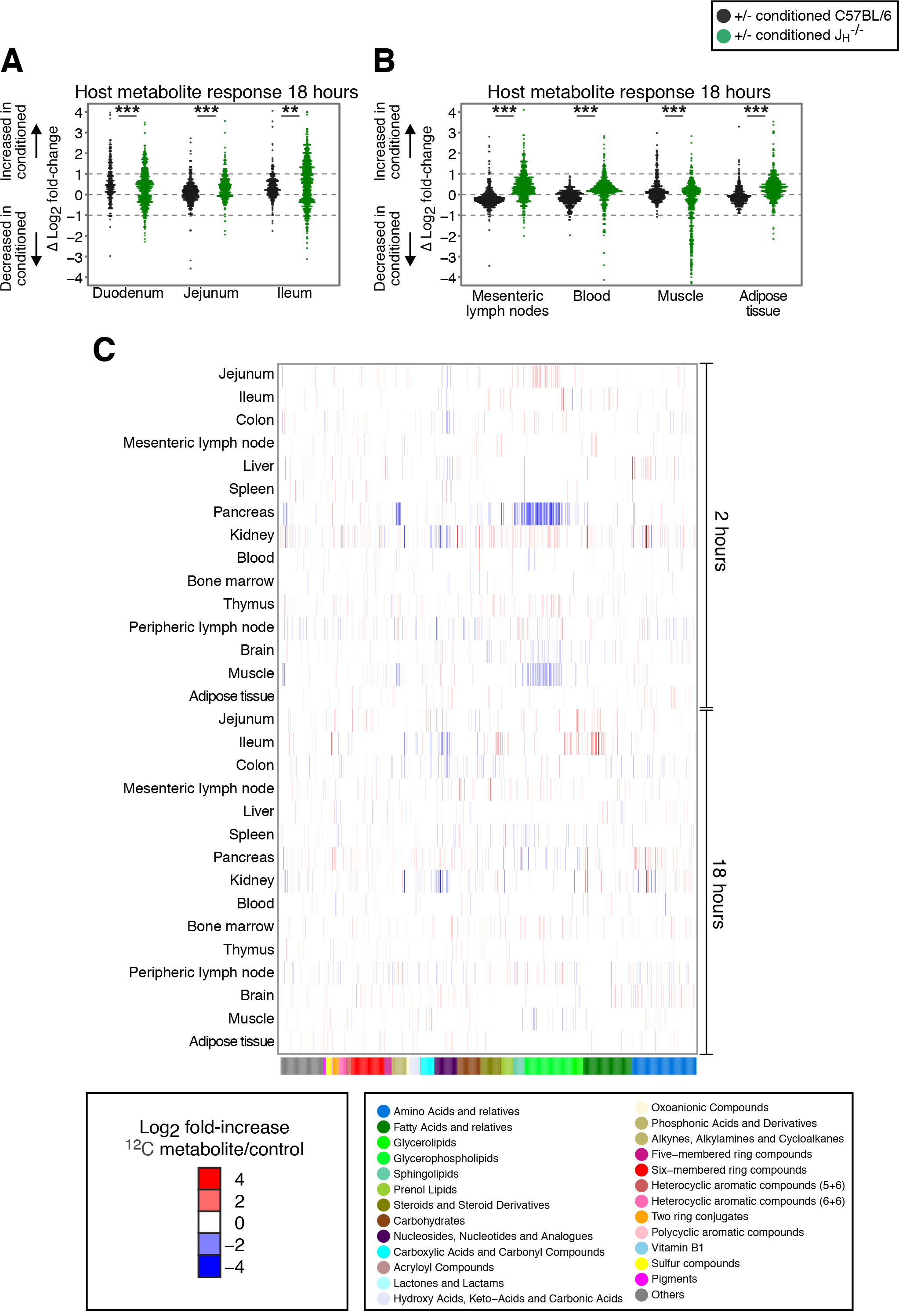
Exaggerated host metabolite responses in antibody-deficient mice following microbial challenge. Germ-free C57BL/6 mice (black symbols) or antibody-deficient J _H_^-/-^ mice (green) were preconditioned with HA107 or control-treated, returning to germ-free status twelve days prior to re-challenge with a gastric dose of fully ^13^C-labelled or ^12^C HA107. Tissue samples were collected after 18 hours and analyzed with Q-TOF mass spectrometry. Annotated ions representing metabolites containing concordant changes of metabolites from ^13^C and ^12^C pulsed mice, without mass shift of the annotated ions were plotted. Log_2_ fold-change differences between preconditioned and control mice for each metabolite is shown for **(A)** duodenal tissue, jejunal tissue and ileal tissue; **(B)** mesenteric lymph nodes, blood, muscle and adipose tissue. **(C)** Heat map showing cluster analysis of concordant increases (red) or losses (blue) of ^12^C-host annotated metabolites (p < 0.05, ≥ 3 mice/group) where the identical metabolite change was seen without a mass shift in both ^13^C- and ^12^C-HA107 challenged antibody deficient mice. Different host tissues at 2 hours and 18 hours after gavage are shown. Metabolite classes are shown in the lower colour bar and legend. Results are representative of two experiments.

**Figure 7.**
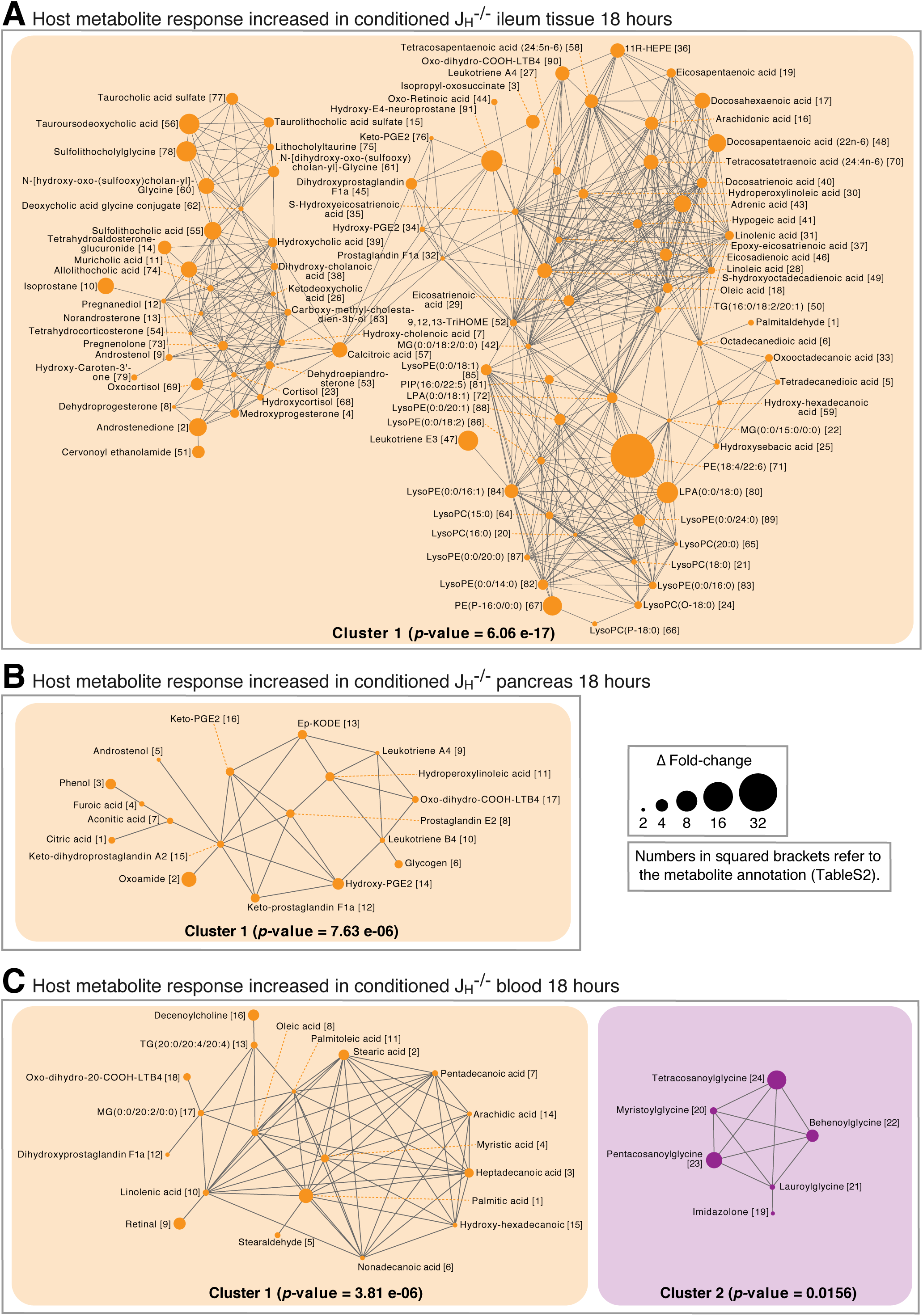
Tanimoto chemical similarity networks of increased metabolite responses in conditioned antibody-deficient mice after microbial challenge. Increased host response metabolites in both ^13^C and ^12^C pulsed mice without mass shift of the annotated ions as in the legend to Figure 6 were applied to Metamapp program to calculate Tanimoto chemical similarity index as described in the legend to Figure 5. The Cytoscape program was used for visualization of networks from the ileum **(A),** the pancreas **(B)** or the blood **(C).** Metabolites with Tanimoto chemical similarity ≥ 0.7 were interconnected. The size of nodes represents the absolute value of difference Δ-fold change between conditioned J _H_^-/-^ and germ-free J _H_^-/-^. Paired Mann-Whitney U test was applied to calculate p-value of each network.

This host metabolomic response was not confined to intestinal tissues. Increased prostaglandins and leukotrienes were also seen in the pancreas of conditioned antibody-deficient mice at 18 hours, following earlier phospholipid breakdown (Figures 6C, S6A and 7B). In the blood of antibody-deficient mice there were also increased host non-esterified fatty acids, as well as glycinated fatty acids (Figure 7C), normally associated with lipid overload through deficient mitochondrial metabolism (Vianey-Liaud et al., 1987). Therefore the lipidemia as a result of bacterial exposure is i) of both bacterial and host origin, and ii) can be triggered by bacterial metabolites in fasting mice by antibody-unprotected exposure to bacteria in the small intestine.

In contrast, the muscle and the spleen of antibody-deficient animals showed a loss of esterified and non-esterified lipids and increased saccharide intermediates (Figures S6B, S7A and S7B). Fat tissue also showed accumulation of saccharide intermediates under these conditions (Figure S7C). These results would be consistent with the established effects of lipotoxicity on lipid and carbohydrate metabolism triggered by circulating non-esterified fatty acids and eicosanoid signalling (Ertunc and Hotamisligil, 2016; Ehrenborg and Krook, 2009).

The host response to increased bacterial dwell-time in the small intestine in the setting of antibody deficiency therefore has both an intestinal and a systemic inflammatory molecular signature, with evidence of systemic lipidemia of both host and bacterial origin, host lipolysis and decreased host glycolysis. These early host molecular changes after a single bacterial challenge dose in antibody-unprotected mice show inflammatory features with the potential to trigger a metabolic syndrome.

## DISCUSSION

The intestinal microbiota is commonly characterised in terms of its composition of different taxa at a single site in the intestine: host responses or a disease phenotype are then correlated with specific taxa or a restricted range of microbial metabolites that are present. Here we present a systematic analysis of the immense extent of microbial-host molecular cross-talk, and show that the microbiota itself induces immune and non-immune boundaries for microbial metabolite penetration into host tissues. A few microbial taxa, such as segmented filamentous bacteria, deliver direct microbial-host signalling through intimate apposition or attachment to the epithelial suface. Yet most of the microbial biomass exists in a highly dynamic system with constant microbial division and death, in the context of continuous transverse and longitudinal flow along the gastrointestinal tract. Here we present an insight into how these dynamics and the immune or non-immune conditioning affect the molecular exchange of both microbial metabolite penetration and the resultant host metabolite responses.

Exposure to intestinal microbes drives a lasting adaptation of the intestinal mucosa through immune and non-immune mechanisms. In the case of humoral immunity, a number of different mechanisms have been described whereby intestinal antibody secretion benefits the host. These include toxin and virus neutralisation, blocking excessive live bacterial translocation, clearance of unwanted macromolecular structures at the epithelial surface and directed sampling of luminal antigen (Kaetzel et al., 1994; Kadaoui and Corthesy, 2007; Macpherson and Uhr, 2004; Lycke et al., 1987; Kaetzel et al., 1991; Burns et al., 1996). Lack of specific IgA has been shown to increase transcriptional evidence of innate immune system activation in the intestine (Peterson et al., 2007). We now show that antibody induction and secretion accelerates intestinal transit and thereby critically limits exposure of the small intestinal mucosa to the pro-inflammatory molecular constituents of intestinal microbes. The transitory colonisation model used generates a dominant and specific secretory IgA response which coats the test organism *in vivo*. We have only found extremely low titres of specific serum IgG after intestinal preconditioning (Gomez de Aguero et al., 2016): our high resolution metabolomic experiments were therefore designed to test the elimination of all antibody isotypes to avoid the compensatory effects of IgM and increased IgG induction in the setting of IgA-deficiency (Harriman et al., 1999), or the enteropathy that can be induced by colonisation in the face of deficiency of the pIgR polymeric immunoglobulin receptor secretory system (Johansen et al., 1999). It is likely that affinity matured secretory IgM can compensate IgA, and a role for IgG isotypes is not excluded.

These effects are poised on the small intestinal mucosa, where limiting bacterial metabolite uptake must be set against the necessity for nutritient absorption. In humans in the developed world the upper small intestine has very low bacterial counts, yet one of the most urgent biomedical problems of our age is to understand why the presence of high densities of small intestinal microbes in children from the undeveloped world causes intestinal dysfunction and infant malnutrition which cannot be adequately treated with caloric increases (Smith et al., 2013a; Kau et al., 2015). Such malnutrition and stunting contributes to the 3 million children that die annually before the age of 5 years and a further 200 million who fail to reach their developmental potential (Grantham-McGregor et al., 2007; Walker et al., 2007). Observations that there are mucosal IgA increases in these children, and IgA is a biomarker of those microbes capable of transferring an enteropathy to germ-free mice, suggests that the mucosal antibody responses are being induced as a protective response in the original host (Beatty et al., 1983; Kau et al., 2015). Limiting microbial metabolite exposure by decreasing the small intestinal dwell time may contribute to the protective effect for those taxa that do not attach to or invade the epithelial surface. Reduction in the intestinal clearance effect is also consistent with the lower small intestinal lymphonodular hyperplasia in the setting of IgA and affinity matured IgM deficiency, owing to a deletion of activation-induced cytidine deaminase (Fagarasan et al., 2002). The host molecular responses are consistent with the observation that individuals affected with environmental enteropathy may paradoxically later be susceptible to the ‘dual burden paradox’ of eventual metabolic syndrome (DeBoer et al., 2012).

We have been able to distinguish bacterial metabolite exposure from the resultant host responses with extremely wide coverage, even in the majority of cases where the same molecule can be potentially derived from either bacterial or host metabolism. The results attest to pervasive microbial metabolite exposure, over many classes of metabolites, including increased bacterial lipid penetration. This effect is even enhanced in the small intestine through transient live bacterial exposure through non immune conditioning effects that include an increase in the intestinal microvascular density. A high proportion of the penetrant microbial metabolites are very rapidly excreted in the urine, although there are also durable increases, some of which benefit the host, such as essential amino acids.

Our model of transitory exposure to live *E.coli* contains some features of the situation in early life humans, when *E.coli* can be prominent and microbiota diversity is limited (Backhed et al., 2015). To be able to interrogate bacterial metabolite penetration at the same time as the host molecular response using stable isotope tracing, the model has to be limited to a single taxon: metabolite exposure from other taxa may vary according to their frequency and metabolic activity in the small intestine. Host responses in the intestine will be shaped by bile acids: although conjugated and deconjugated bile acids are present in our system, the specifics of subsequent dihydroxylation and epimerisation of the cyclopentaphenanthrene ring substitutions depend on the diversity of the microbiota, which was here minimised (Macpherson et al., 2016). We did not find evidence that the cell wall auxotrophy of HA107 increased the fragility of the organism compared with its non-auxotrophic parent strain, and we could grow viable bacteria from fecal samples, provided the auxotrophic requirements were added to the culture medium. Although challenge HA107 bacteria in these experiments could be efficiently recovered from feces, bacterial death in the mouse intestine contributes to the metabolite load. This is not restricted to the HA107 model. By measuring *in vivo* replication rates and overall dynamics, both we and others have calculated high rates of bacterial death in the intestine in steady state monocolonised animals, amounting to 1 in 8 divisions of Escherichia coli *in vivo* (Hai et al., 2015; Myhrvold et al., 2015). Through reducing the bacterial dwell time in the small intestine, antibodies likely lessen its exposure to metabolites from dead bacteria and subsequent uptake into the body.

Whilst the breadth of metabolic exchange allows a broad portfolio of metabolic responses, the predominant effect in antibody-deficient mice was lipidemia of both microbial and host origin, and host lipid breakdown in the intestine and the muscle. Breakdown of phospholipid components in the intestine also is accompanied by prostaglandin and leukotriene synthesis. This picture, including the existence of host aminoacetylated fatty acids in the serum and presence of intermediates of gluconeogenesis indicates likely lipotoxicity and insulin resistance (Ertunc and Hotamisligil, 2016; Ehrenborg and Krook, 2009). Increased accumulation of body fat in non-inflamed immune-competent colonised animals has been established as a result of microbial modulated angiogenin-4 and fasting-induced adipose factor signalling, supplemented by an increased energy yield from harvesting of resistant dietary carbohydrates (Backhed et al., 2004; Backhed et al., 2007). Lipidemia of a combined microbial and host metabolite origin can now be demonstrated by challenging the intestine with microbes in the absence of antibody protection: this likely contributes to a range of clinically relevant disorders in the presence of a ‘dysbiotic’ microbiota (or dysfunctional control of the microbes present), including non-alcoholic fatty liver disease (Boursier et al., 2016) and cardiovascular disease (Wang et al., 2011). Extending our model could potentially separate host or dietary molecular effects from microbial molecular effects in understanding these disorders.

A substantial investment of metabolic energy is made in the secretory IgA response which comprises approximately 75% of all immunoglobulin production in mammals (Macpherson et al., 2008), most of which is secreted in the small intestine. The small intestine is a delicately positioned interface with the environment. On one hand, it is specialised for uptake of nutrients with minimal mucus covering and a combination of digestive and secretory mechanisms. On the other, it must handle the inevitable contamination with intestinal microbes: this it does with a combination of physiological adaptations to clear microbial compounds quickly into the urine and secretory immunoglobulins accelerating transit away from the susceptible small intestinal mucosa, thereby modulating both microbial penetration into the host and the subsequent host response.

## AUTHOR CONTRIBUTIONS

A.J.M. and U.S. conceived and designed the project. A.J.M., U.S., and K.D.M. supervised the experiments. Y.U., H.L., M.A.L., M.G.A., B.Y., F.R., S.C.G. performed metabolomics sampling. T.F. and M.Z. performed mass spectrometric measurement and data processing. Y.U., T.F. and A.J.M. performed the bioinformatics analysis. H.L. and M.A.L. performed the intestinal bacterial transit experiments in Figure 4C and 4D. M.S. assisted with live intestinal endoscopy experiments in Figure 3E. S.H. provided the E. coli strain HA107 and contributed to validation of its use for this project. A.J.M., U.S., Y.U. and T.F. wrote the manuscript.

## ACKNOWLEDGEMENTS

We thank the staff of the clean mouse facility of the University of Bern and Professors Wolf Hardt and Christoph Mueller for their helpful comments and input. We thank Dr. Irene Keller for the processing of RNAseq data and Professor D. Candinas and the Genaxen Foundation for support with the Clean Mouse Facility. The work was supported by the Swiss National Science foundation (SNSF 310030B_160262, SNF Sinergia CRSII3_136286 to AJM and SNSF Sinergia CRSII3_154414 AJM and US) and the Systems X program (GutX AJM and US).

## LIST OF SUPPLEMENTARY FIGURES

Figure S1 – related to Figure 1

Figure S2 – related to Figure 2

Figure S3 – related to Figure 3

Figure S4 – related to Figure 4

Figure S5 – related to Figure 5

Figure S6 – related to Figures 6 and 7

Figure S7 – related to Figure 7

## STAR METHODS

### Mice

Germ-free C57BL/6(J) and J _H_^-/-^ (Chen et al., 1993) mice were born and maintained in flexible-film isolators in the Clean Mouse Facility, University of Bern, Switzerland. Age and gender-matched mice were used at 10-12 weeks for the metabolomics samplings in this study. All mice were independently confirmed to be pathogen-free. They were also verified germ-free within the breeding isolators and during experiments (apart from the transient colonization and the colonized experiment shown in Figures S5B and S5C) by culture-dependent and -independent methods. All mouse experiments were performed in accordance with Swiss Federal and Cantonal regulations.

### Preconditioning the intestinal mucosa through transient bacterial colonisation in germ-free mice

*Escherichia coli* HA107 was cultured in LB medium supplemented with 100 μg/ml *meso*-DAP and 400 μg/ml **D**-alanine (Hapfelmeier et al., 2010). Bacterial cultures were centrifuged at 3480**g** for 10 min at 4°C. After washing twice with sterile PBS, the required dose of bacteria was resuspended in PBS (10^10^ CFU per 500 μl). The bacterial solution was gavaged directly into the mouse stomach on alternate days four times. One week after the last HA107 gavage, the feces from the mice were routinely verified by culture-dependent and -independent methods to be germ-free. Control germ-free mice were similarly treated in parallel throughout, although gavaged with an equivalent volume of sterile PBS. Preconditioned mice were studied further 14 days after the last conditioning dose.

### Preparation of ^13^C-labelled bacteria

To prepare ^13^C-labelled HA107, sterile ^13^C-labelled bacterial extracts from *E. coli* MG1655 were used as source of ^13^C-labelled amino acids and vitamins. For this, *E. coli* MG1655 was initially cultured in M9 medium using ^13^C-**D**-glucose (Cambridge Isotope Laboratories) as sole carbon source. After three sequential subcultures, fully ^13^C-labelled MG1655 bacterial extracts were obtained and sterilized by filtration (0.22 μm). *E. coli* HA107 was then cultured in M9 medium containing ^13^C-**D**-glucose and ^13^C-labelled bacterial extracts from *E. coli* MG1655. After three sequential cultures, the HA107 bacterial culture was centrifuged at 3480 **g** for 10 min at 4°C. After washing twice with sterile PBS, the required dose of HA107 bacteria was suspended in PBS (10^10^ CFU per 500 μl). We verified that this protocol achieved essentially complete (> 90%) ^13^C-labelling of the carbon chains of all bacterial molecules across all compound classes (Figures S1D and S1E).

### Mass spectrometry

Samples were snap-frozen in liquid nitrogen. To prepare fluid samples for mass spectrometry, weighed fluid samples (50 g) were precipitated with final 80% methanol for 1 hour at -20°C. After centrifugation at 20,000 **g** for 10 min at room temperature, the supernatant was transferred to Q-TOF plates. To prepare tissue samples for mass spectrometry, metabolites from weighed tissue samples (50 g) were extracted with 20 times weight of Millipore water (80°C) followed by homogenization at 25 Hz for 2 min in a tissue homogenizer (Retsch MM 400) with a clean steel ball (Berani). After centrifugation at 20,000 **g** for 10 min at room temperature, supernatants were transferred to Q-TOF plates.

Quantification of relative metabolites levels was carried out with an Agilent 6550 Q-TOF mass spectrometer (Agilent Technologies Inc., Santa Clara, CA) by non-targeted flow injection analysis as described previously (Fuhrer et al 2011). Profile spectra with high mass accuracy were recorded from 50 to 1000 m/z in negative ionization mode. The common mass axis after sample alignment was recalibrated using known frequently occuring metabolite ions. Ions were finally annotated based on accurate mass comparison using 1 mDa mass tolerance against 9261 unique metabolites present in a pooled list of compounds derived from the Human Metabolome Database (Wishart et al., 2013), *E. coli* genome scale-model (Orth et al., 2011) and the E. coli Metabolome Database (Sajed et al., 2016). All potentially possible ^13^C isotopes were annotated taking into acount the respective sum mass formula.

### Determining the extent of ^13^C-labelling in *E. coli* HA107 used for stable isotope tracing

The free intracellular metabolites were extracted from the cell pellet with Millipore water at 80°C and analyzed by non-targeted flow injection analysis as described above. Proteinogenic amino acids were obtained from the extracted pellet after hydrolysis with 6 M HCl and subsequently derivatized with N-methyl-N-(tert-butyldimethylsilyl)-trifluoroacetamide as described previously (Nanchen et al., 2007). The ^13^C-labelling patterns were then determined on a 6890N Network GC system with a 5975 inert XL mass selective detector (Agilent Technologies Inc., Santa Clara, CA). Fractional label for both intracellular metabolites and proteinogenic amino acids was then estimated from the relative abundances of m_0_ and m_max_ ion counts as m_max_ / (m_0_ + m_max_).

### Metabolomics data analysis

Log fold-changes of ^13^C-labelled compounds were calculated from the division of [positive ion counts at m+x isotope annotation from ^13^C-HA107 gavaged samples] by [ion counts at m+x isotope annotation from PBS gavaged samples]. ?Log_2_ fold-change was calculated from the subtraction of [Log_2_ fold-change of germ-free control mice] from [Log fold-change of conditioned mice]. Log fold-changes of ^12^C compounds in ^13^C-HA107 gavaged samples were calculated from the division of [ion counts at m annotation from ^13^C-HA107 gavaged samples] by [ion counts at m_0_ annotation from PBS gavaged samples]. In the same way, Log fold-change of ^12^C compounds in ^12^C-HA107 gavaged samples were calculated from the division of [ion counts at m annotation from ^12^C-HA107 gavaged samples] by [ion counts at m annotation from PBS gavaged samples]. For the host response of endogenous ^12^C compounds, compounds with the same direction of Log fold-change in both Log fold-change of ^12^C compounds in ^13^C-HA107 gavaged samples and Log_2_ fold-change of ^12^C compounds in ^12^C-HA107 gavaged samples were taken. Where shown, the Metamapp program was used to evaluate the Tanimoto chemical similarity index on compounds selectively present in one condition applying Cytoscape software for network visualization. Mann Whitney U test was performed using wilcox.test function in R (* = p < 0.05, ** = p < 0.01, *** = p < 0.001).

### Preparation of ^14^C-labelled bacteria

M9 medium supplemented with 100 μg/ml meso-DAP, 400 μg/ml D-alanine, 10 μg/ml lysine, 10 μg/ml threonine, 10 μg/ml methionine, 10 μg/ml proline, 4 μg/ml thiamine, 2 mM L-arabinose, and 5 mM succinate was inoculated with 1/100^th^ amount of *E. coli* HA107 (or JM83) overnight culture grown in LB medium. The culture was incubated with shaking at 180 rpm at 37 °C. When the culture reached OD_600_ 0.28±0.05, 2-[1-^14^C]-deoxy-D-glucose (Perkin Elmer) was added. The bacterial culture was further incubated for 12-14 hours at 37 °C. After harvesting the bacterial culture, the bacterial pellet was washed twice with PBS to remove unincorporated ^14^C-radioactive materials and resuspended prior to gavage.

### Cecal injection

For cecal injection mice were anaesthetized with isoflurane by inhalation (3.0 to 4.0% v/v isoflurane from a vapourizer with O_2_ flow at 1-2 L/min). An incision (∼ 1 cm) was made with curved scissors at left side of lower abdomen, and the cecum was located. ^14^C-labelled HA107 was injected into the middle of cecum and mixed thoroughly with the contents. The abdominal incision was closed with AUTOCLIP Wound Clip System (BD). Temgesic (0.05-0.1 μg/g body weight) was injected subcutaneously for analgesia.

### Liquid scintillation counting

Weighed samples were incubated with 1 ml of NCS Tissue Solubilizer (GE Healthcare) at 56°C until completely dissolved. The pH was neutralized with 200 μl of 100% acetic acid. Ultima Gold liquid scintillation cocktail (18 ml, PerkinElmer) was added to each sample. The ^14^C-radioactivity in each sample was measured in a TRI-CARB 2300TR Liquid Scintillation Analyzer (Packard). Background scintillation levels for each fluid or tissue type were subtracted using matched samples from control germ-free mice that had not received ^14^C-labelled bacteria.

### RNA isolation

Tissue samples were snap-frozen in TRIzol reagent (Invitrogen) using liquid nitrogen. Thawed tissues in TRIzol reagent were immediately homogenized at 25 Hz for 2 min in a tissue homogenizer (Retsch MM 400) with sterile steel balls (Berani). Total RNA was isolated following the standard protocol from Invitrogen. The total RNA was further purified with RNeasy MinElute kit (Qiagen). RNA quality was assessed on a Bioanalyzer 2100 (Agilent). Ribosomal RNA was removed with Ribominus Transcriptome Isolation Kit (Invitrogen). RNA libraries were prepared with TruSeq RNA sample preparation v2 kit (TruSeq Stranded mRNA Sample Preparation, Illumina). RNA sequencing was performed with Illumina HiSeq 2500 using the 150bp paired-end mode. Read quality control was with FastQC. Each read was mapped onto the mouse genome using TopHat2. Counts per gene were assessed using HTSeq-count software. Differential gene expression analysis was performed using DESeq2. Adjusted Log_2_ fold-change and adjusted p-values (corrected for multiple testing) were used to determine significant differences in gene expression. Functional protein association networks of genes of adjusted p-value of < 0.05 were analyzed using the STRING database (version 10) taking network confidence scores ≥ 0.4.

### Live intestinal endomicroscopy

Mice investigated with endomicroscopy were anaesthetized with isoflurane inhalation as described for cecal injection on a heating pad at 37°C. The abdomen of the anaesthetized mouse was opened and a Cellvizio confocal miniprobe (Mauna Kea) was inserted into the intestinal ileal lumen. Fluorescein isothiocyanate (FITC)-dextran (0.5 mg, 70 kDa, Sigma) in PBS was injected intravenously to provide real-time vascular contrast. The fluorescence of FITC in the intestinal vasculature was recorded with Cellvizio software (Mauna Kea). Metrics of the endothelial and epithelial layers were evaluated with Image J software (NIH).

### Enzyme-linked immunosorbent assay (ELISA)

The gastrointestinal tracts from HA107 conditioned germ-free C57BL/6 mice were separated according to section (duodenum, jejunum, cecum and colon). Isolated luminal contents from each section were weighed. IgA concentration was measured with ELISA using goat anti-mouse IgA (Bio-Rad Antibodies) coating antibody. The detection antibody was HRP-conjugated anti-mouse IgA (Sigma). Standards were myeloma-derived purified IgA (Hycult). The IgA concentration in the contents of each germ-free intestinal segment was plotted.

### Bacterial immunofluorescent staining

*E. coli* HA107 was cultured without shaking at 37°C for overnight. The bacteria were then washed in PBS and resuspended at 10^7^ per ml. Portions (50 μl) of the bacterial suspension were dropped onto glass slides and the bacterial drop was encircled using a wax pen. Bacteria were allowed to sediment onto the glass slides and to dry. Transiently flamed bacteria on glass slides were stained with intestinal wash prepared from small intestine of HA107 conditioned germ-free C57BL/6 mice. The IgA antibodies bound to bacteria were visualized with FITC-conjugated anti-mouse IgA (BD Bioscience). DAPI (4’, 6-diamidino-2-phenylindole) was used to counterstain the bacterial DNA. Immunofluorescent pictures were taken using Flash Slide Scanner (PerkinElmer) with 40 magnifications.

### Bacterial intestinal transit time

Germ-free C57BL/6 and J _H_^-/-^ mice were conditioned with HA107 as described previously. For small intestinal transit, 10^7^ CFU of HA107 were introduced directly into the stomachs of mice. Two hours after the intragastric gavage, small intestines were collected and separated into 10 sections of identical length from duodenum to ileum. All luminal contents in each section were plated on LB agar plates with the auxotrophic supplements and results were evaluated for each segment. To measure whole intestinal transit time, 10^10^ CFU of HA107 were introduced to the stomachs of mice. Feces were sequentially collected and cultured as described previously.

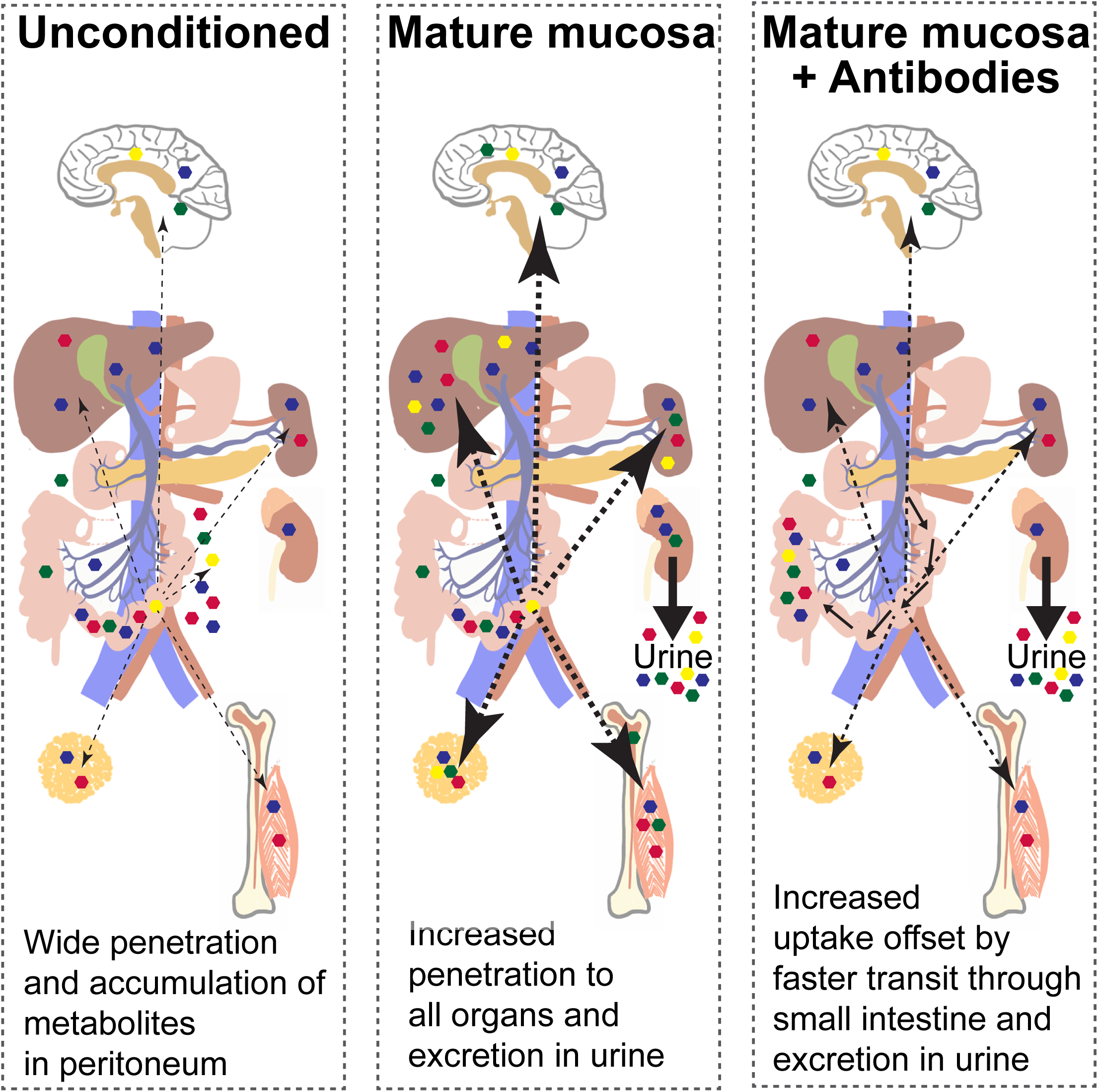

## References

Adamson, R.H., Bridges, J.W., Evans, M.E., and Williams, R.T. (1970). Species differences in the aromatization of quinic acid in vivo and the role of gut bacteria. Biochem J 116, 437–443.

Arpaia, N., Campbell, C., Fan, X., Dikiy, S., van der Veeken, J., deRoos, P., Liu, H., Cross, J.R., Pfeffer, K., Coffer, P.J., et al. (2013). Metabolites produced by commensal bacteria promote peripheral regulatory T-cell generation. Nature 504, 451–455.

Backhed, F., Ding, H., Wang, T., Hooper, L.V., Koh, G.Y., Nagy, A., Semenkovich, C.F., and Gordon, J.I. (2004). The gut microbiota as an environmental factor that regulates fat storage. Proc Natl Acad Sci U S A 101, 15718–15723.

Backhed, F., Manchester, J.K., Semenkovich, C.F., and Gordon, J.I. (2007). Mechanisms underlying the resistance to diet-induced obesity in germ-free mice. Proc Natl Acad Sci U S A 104, 979–984.

Backhed, F., Roswall, J., Peng, Y., Feng, Q., Jia, H., Kovatcheva-Datchary, P., Li, Y., Xia, Y., Xie, H., Zhong, H., et al. (2015). Dynamics and Stabilization of the Human Gut Microbiome during the First Year of Life. Cell Host Microbe 17, 690–703.

Beatty, D.W., Napier, B., Sinclair-Smith, C.C., McCabe, K., and Hughes, E.J. (1983). Secretory IgA synthesis in Kwashiorkor. J Clin Lab Immunol 12, 31–36.

Belkaid, Y., and Harrison, O.J. (2017). Homeostatic Immunity and the Microbiota. Immunity 46, 562–576.

Blumberg, R., and Powrie, F. (2012). Microbiota, disease, and back to health: a metastable journey. Sci Transl Med 4, 137rv137.

Boursier, J., Mueller, O., Barret, M., Machado, M., Fizanne, L., Araujo-Perez, F., Guy, C.D., Seed, P.C., Rawls, J.F., David, L.A., et al. (2016). The severity of nonalcoholic fatty liver disease is associated with gut dysbiosis and shift in the metabolic function of the gut microbiota. Hepatology 63, 764–775.

Brugiroux, S., Beutler, M., Pfann, C., Garzetti, D., Ruscheweyh, H.J., Ring, D., Diehl, M., Herp, S., Lotscher, Y., Hussain, S., et al. (2016). Genome-guided design of a defined mouse microbiota that confers colonization resistance against Salmonella enterica serovar Typhimurium. Nat Microbiol 2, 16215.

Bunker, J.J., Flynn, T.M., Koval, J.C., Shaw, D.G., Meisel, M., McDonald, B.D., Ishizuka, I.E., Dent, A.L., Wilson, P.C., Jabri, B., et al. (2015). Innate and Adaptive Humoral Responses Coat Distinct Commensal Bacteria with Immunoglobulin A. Immunity 43, 541–553.

Burns, J.W., Siadat-Pajouh, M., Krishnaney, A.A., and Greenberg, H.B. (1996). Protective effect of rotavirus VP6-specific IgA monoclonal antibodies that lack neutralizing activity. Science 272, 104–107.

Chen, J.Z., Trounstine, M., Alt, F.W., Young, F., Kurahara, C., Loring, J.F., and Huszar, D. (1993). Immunoglobulin Gene Rearrangement in B-Cell Deficient Mice Generated By Targeted Deletion of the J(H) Locus. International Immunology 5, 647–656.

Cotran, R., Kendrick, M.I., and Kass, E.H. (1960). Role of intestinal bacteria in aromatization of quinic acid in man and guinea pig. Proc Soc Exp Biol Med 104, 424–426.

Dayman, J., and Jepson, J.B. (1969). The metabolism of caffeic acid in humans: the dehydroxylating action of intestinal bacteria. Biochem J 113, 11P.

de Graaf, A.A., Maathuis, A., de Waard, P., Deutz, N.E., Dijkema, C., de Vos, W.M., and Venema, K. (2010). Profiling human gut bacterial metabolism and its kinetics using [U-13C]glucose and NMR. NMR Biomed 23, 2–12.

DeBoer, M.D., Lima, A.A., Oria, R.B., Scharf, R.J., Moore, S.R., Luna, M.A., and Guerrant, R.L. (2012). Early childhood growth failure and the developmental origins of adult disease: do enteric infections and malnutrition increase risk for the metabolic syndrome? Nutr Rev 70, 642–653.

Donowitz, J.R., Haque, R., Kirkpatrick, B.D., Alam, M., Lu, M., Kabir, M., Kakon, S.H., Islam, B.Z., Afreen, S., Musa, A., et al. (2016). Small Intestine Bacterial Overgrowth and Environmental Enteropathy in Bangladeshi Children. MBio 7, e02102–02115.

Ehrenborg, E., and Krook, A. (2009). Regulation of skeletal muscle physiology and metabolism by peroxisome proliferator-activated receptor delta. Pharmacol Rev 61, 373–393.

El Aidy, S., Merrifield, C.A., Derrien, M., van Baarlen, P., Hooiveld, G., Levenez, F., Dore, J., Dekker, J., Holmes, E., Claus, S.P., et al. (2013). The gut microbiota elicits a profound metabolic reorientation in the mouse jejunal mucosa during conventionalisation. Gut 62, 1306–1314.

Ertunc, M.E., and Hotamisligil, G.S. (2016). Lipid signaling and lipotoxicity in metaflammation: indications for metabolic disease pathogenesis and treatment. J Lipid Res 57, 2099–2114.

Fagarasan, S., Muramatsu, M., Suzuki, K., Nagaoka, H., Hiai, H., and Honjo, T. (2002). Critical roles of activation-induced cytidine deaminase in the homeostasis of gut flora. Science 298, 1424–1427.

Fuhrer, T., Heer, D., Begemann, B., and Zamboni, N. (2011). High-throughput, accurate mass metabolome profiling of cellular extracts by flow injection-time-of-flight mass spectrometry. Anal Chem 83, 7074-7080.

Fuhrer, T., Zampieri, M., Sevin, D.C., Sauer, U., and Zamboni, N. (2017). Genomewide landscape of gene-metabolome associations in Escherichia coli. Mol Syst Biol 13, 907.

Gensollen, T., Iyer, S.S., Kasper, D.L., and Blumberg, R.S. (2016). How colonization by microbiota in early life shapes the immune system. Science 352, 539–544.

Gomez de Aguero, M., Ganal-Vonarburg, S.C., Fuhrer, T., Rupp, S., Uchimura, Y., Li, H., Steinert, A., Heikenwalder, M., Hapfelmeier, S., Sauer, U., et al. (2016). The maternal microbiota drives early postnatal innate immune development. Science 351, 1296–1302.

Grantham-McGregor, S., Cheung, Y.B., Cueto, S., Glewwe, P., Richter, L., Strupp, B., and International Child Development Steering, G. (2007). Developmental potential in the first 5 years for children in developing countries. Lancet 369, 60–70.

Hai, L., Limenitakis, J., Fuhrer, T., Geuking, M.B., Lawson, M.A., Wyss, M., Brugiroux, S., Keller, I., Macpherson, J.A., Rupp, S., et al. (2015). The outer mucus layer hosts a distinct intestinal microbial niche. Nature Communications 6, 8292.

Hapfelmeier, S., Lawson, M.A., Slack, E., Kirundi, J.K., Stoel, M., Heikenwalder, M., Cahenzli, J., Velykoredko, Y., Balmer, M.L., Endt, K., et al. (2010). Reversible microbial colonization of germ-free mice reveals the dynamics of IgA immune responses. Science 328, 1705–1709.

Harriman, G.R., Bogue, M., Rogers, P., Finegold, M., Pacheco, S., Bradley, A., Zhang, Y., and Mbawuike, I.N. (1999). Targeted deletion of the IgA constant region in mice leads to IgA deficiency with alterations in expression of other Ig isotypes. J Immunol 162, 2521–2529.

Hooper, L.V., Wong, M.H., Thelin, A., Hansson, L., Falk, P.G., and Gordon, J.I. (2001). Molecular analysis of commensal host-microbial relationships in the intestine. Science 291, 881–884.

Johansen, F.E., Pekna, M., Norderhaug, I.N., Haneberg, B., Hietala, M.A., Krajci, P., Betsholtz, C., and Brandtzaeg, P. (1999). Absence of epithelial immunoglobulin A transport, with increased mucosal leakiness, in polymeric immunoglobulin Receptor/Secretory component-deficient mice. J Exp Med 190, 915–922.

Johansson, M.E., Phillipson, M., Petersson, J., Velcich, A., Holm, L., and Hansson, G.C. (2008). The inner of the two Muc2 mucin-dependent mucus layers in colon is devoid of bacteria. Proc Natl Acad Sci U S A 105, 15064–15069.

Jones, H.F., Davidson, G.P., Brooks, D.A., and Butler, R.N. (2011). Is small-bowel bacterial overgrowth an underdiagnosed disorder in children with gastrointestinal symptoms? J Pediatr Gastroenterol Nutr 52, 632–634.

Kadaoui, K.A., and Corthesy, B. (2007). Secretory IgA mediates bacterial translocation to dendritic cells in mouse Peyer’s patches with restriction to mucosal compartment. J Immunol 179, 7751–7757.

Kaetzel, C.S., Robinson, J.K., Chintalacharuvu, K.R., Vaerman, J.P., and Lamm, M.E. (1991). The polymeric immunoglobulin receptor (secretory component) mediates transport of immune complexes across epithelial cells: a local defense function for IgA. Proc Natl Acad Sci U S A 88, 8796–8800.

Kaetzel, C.S., Robinson, J.K., and Lamm, M.E. (1994). Epithelial transcytosis of monomeric IgA and IgG cross-linked through antigen to polymeric IgA. A role for monomeric antibodies in the mucosal immune system. J Immunol 152, 72–76.

Kau, A.L., Planer, J.D., Liu, J., Rao, S., Yatsunenko, T., Trehan, I., Manary, M.J., Liu, T.C., Stappenbeck, T.S., Maleta, K.M., et al. (2015). Functional characterization of IgA-targeted bacterial taxa from undernourished Malawian children that produce diet-dependent enteropathy. Sci Transl Med 7, 276ra224.

Korpe, P.S., and Petri, W.A. Jr. (2012). Environmental enteropathy: critical implications of a poorly understood condition. Trends Mol Med 18, 328–336.

Leslie, C.C. (2015). Cytosolic phospholipase A(2): physiological function and role in disease. J Lipid Res 56, 1386–1402.

Lycke, N., Eriksen, L., and Holmgren, J. (1987). Protection against cholera toxin after oral immunisation is thymus dependent and associated with intestinal production of neutralising IgA antitoxin. Scand. J. Immunol. 25, 413–419.

Macpherson, A.J., Gatto, D., Sainsbury, E., Harriman, G.R., Hengartner, H., and Zinkernagel, R.M. (2000). A primitive T cell-independent mechanism of intestinal mucosal IgA responses to commensal bacteria. Science 288, 2222–2226.

Macpherson, A.J., Heikenwalder, M., and Ganal-Vonarburg, S.C. (2016). The Liver at the Nexus of Host-Microbial Interactions. Cell Host Microbe 20, 561–571.

Macpherson, A.J., McCoy, K.D., Johansen, F.E., and Brandtzaeg, P. (2008). The immune geography of IgA induction and function. Mucosal Immunol 1, 11–22.

Macpherson, A.J., and Uhr, T. (2004). Induction of protective IgA by intestinal dendritic cells carrying commensal bacteria. Science 303, 1662–1665.

Myhrvold, C., Kotula, J.W., Hicks, W.M., Conway, N.J., and Silver, P.A. (2015). A distributed cell division counter reveals growth dynamics in the gut microbiota. Nat Commun 6, 10039.

Nanchen, A., Fuhrer, T., and Sauer, U. (2007). Metabolomics methods and protocols. In Methods in Molecular Biology. W. Weckworth, ed. (Humana Press, Springer), pp. 177–197.

Nicholls, A.W., Mortishire-Smith, R.J., and Nicholson, J.K. (2003). NMR spectroscopic-based metabonomic studies of urinary metabolite variation in acclimatizing germ-free rats. Chem Res Toxicol 16, 1395–1404.

Orth, J.D., Conrad, T.M., Na, J., Lerman, J.A., Nam, H., Feist, A.M., and Palsson, B.O. (2011). A comprehensive genome-scale reconstruction of Escherichia coli metabolism-2011. Mol Syst Biol 7, 535.

Padmanabhan, P., Grosse, J., Asad, A.B., Radda, G.K., and Golay, X. (2013). Gastrointestinal transit measurements in mice with 99mTc-DTPA-labeled activated charcoal using NanoSPECT-CT. EJNMMI Res 3, 60.

Peppercorn, M.A., and Goldman, P. (1972). Caffeic acid metabolism by gnotobiotic rats and their intestinal bacteria. Proc Natl Acad Sci U S A 69, 1413–1415.

Perry, R.J., Peng, L., Barry, N.A., Cline, G.W., Zhang, D., Cardone, R.L., Petersen, K.F., Kibbey, R.G., Goodman, A.L., and Shulman, G.I. (2016). Acetate mediates a microbiome-brain-beta-cell axis to promote metabolic syndrome. Nature 534, 213–217.

Peterson, D.A., McNulty, N.P., Guruge, J.L., and Gordon, J.I. (2007). IgA response to symbiotic bacteria as a mediator of gut homeostasis. Cell Host Microbe 2, 328–339.

Petersson, J., Schreiber, O., Hansson, G.C., Gendler, S.J., Velcich, A., Lundberg, J.O., Roos, S., Holm, L., and Phillipson, M. (2011). Importance and regulation of the colonic mucus barrier in a mouse model of colitis. Am J Physiol Gastrointest Liver Physiol 300, G327–333.

Sajed, T., Marcu, A., Ramirez, M., Pon, A., Guo, A.C., Knox, C., Wilson, M., Grant, J.R., Djoumbou, Y., and Wishart, D.S. (2016). ECMDB 2.0: A richer resource for understanding the biochemistry of E. coli. Nucleic Acids Res 44, D495–501.

Salzman, N.H., Hung, K., Haribhai, D., Chu, H., Karlsson-Sjoberg, J., Amir, E., Teggatz, P., Barman, M., Hayward, M., Eastwood, D., et al. (2010). Enteric defensins are essential regulators of intestinal microbial ecology. Nat Immunol 11, 76–83.

Scheline, R.R., and Midtvedt, T. (1970). Absence of dehydroxylation of caffeic acid in germ-free rats. Experientia 26, 1068–1069.

Sevin, D.C., Fuhrer, T., Zamboni, N., and Sauer, U. (2017). Nontargeted in vitro metabolomics for high-throughput identification of novel enzymes in Escherichia coli. Nat Methods 14, 187–194.

Smith, K., McCoy, K.D., and Macpherson, A.J. (2007). Use of axenic animals in studying the adaptation of mammals to their commensal intestinal microbiota. Semin Immunol 19, 59–69.

Smith, M.I., Yatsunenko, T., Manary, M.J., Trehan, I., Mkakosya, R., Cheng, J., Kau, A.L., Rich, S.S., Concannon, P., Mychaleckyj, J.C., et al. (2013a). Gut microbiomes of Malawian twin pairs discordant for kwashiorkor. Science 339, 548–554.

Smith, P.M., Howitt, M.R., Panikov, N., Michaud, M., Gallini, C.A., Bohlooly, Y.M., Glickman, J.N., and Garrett, W.S. (2013b). The microbial metabolites, short-chain fatty acids, regulate colonic Treg cell homeostasis. Science 341, 569–573.

Sommer, F., and Backhed, F. (2013). The gut microbiota-masters of host development and physiology. Nat Rev Microbiol 11, 227–238.

Stappenbeck, T.S., Hooper, L.V., and Gordon, J.I. (2002). Developmental regulation of intestinal angiogenesis by indigenous microbes via Paneth cells. Proc Natl Acad Sci U S A 99, 15451–15455.

Tannock, G.W., Lawley, B., Munro, K., Sims, I.M., Lee, J., Butts, C.A., and Roy, N. (2014). RNA-stable-isotope probing shows utilization of carbon from inulin by specific bacterial populations in the rat large bowel. Appl Environ Microbiol 80, 2240–2247.

Vaishnava, S., Yamamoto, M., Severson, K.M., Ruhn, K.A., Yu, X., Koren, O., Ley, R., Wakeland, E.K., and Hooper, L.V. (2011). The antibacterial lectin RegIIIgamma promotes the spatial segregation of microbiota and host in the intestine. Science 334, 255–258.

Vianey-Liaud, C., Divry, P., Gregersen, N., and Mathieu, M. (1987). The inborn errors of mitochondrial fatty acid oxidation. J Inherit Metab Dis 10 Suppl 1, 159–200.

Walker, S.P., Wachs, T.D., Gardner, J.M., Lozoff, B., Wasserman, G.A., Pollitt, E., Carter, J.A., and International Child Development Steering, G. (2007). Child development: risk factors for adverse outcomes in developing countries. Lancet 369, 145–157.

Wang, Z., Klipfell, E., Bennett, B.J., Koeth, R., Levison, B.S., Dugar, B., Feldstein, A.E., Britt, E.B., Fu, X., Chung, Y.M., et al. (2011). Gut flora metabolism of phosphatidylcholine promotes cardiovascular disease. Nature 472, 57–63.

Wikoff, W.R., Anfora, A.T., Liu, J., Schultz, P.G., Lesley, S.A., Peters, E.C., and Siuzdak, G. (2009). Metabolomics analysis reveals large effects of gut microflora on mammalian blood metabolites. Proc Natl Acad Sci U S A 106, 3698–3703.

Williams, R.E., Eyton-Jones, H.W., Farnworth, M.J., Gallagher, R., and Provan, W.M. (2002). Effect of intestinal microflora on the urinary metabolic profile of rats: a (1)H-nuclear magnetic resonance spectroscopy study. Xenobiotica 32, 783–794.

Wishart, D.S., Jewison, T., Guo, A.C., Wilson, M., Knox, C., Liu, Y., Djoumbou, Y., Mandal, R., Aziat, F., Dong, E., et al. (2013). HMDB 3.0-The Human Metabolome Database in 2013. Nucleic Acids Res 41, D801–807.

